# Niche-targeted therapy via YAP/TAZ activation enhances hematopoietic regeneration

**DOI:** 10.1101/2025.08.21.671455

**Authors:** Shun Uemura, Masayuki Yamashita, Takako Yokomizo-Nakano, Ayako Aihara, Takumi Iwawaki, Shuhei Koide, Yaeko Nakajima-Takagi, Motohiko Oshima, Yoshiki Omatsu, Bahityar Rahmutulla, Atsushi Kaneda, Miki Nishio, Akira Suzuki, Takashi Nagasawa, Kenta Kagaya, Taito Nishino, Atsushi Iwama

**Author notes:** **Corresponding authors:** Atsushi Iwama, M.D. Ph.D., Division of Stem Cell and Molecular Medicine, Center for Stem Cell Biology and Regenerative Medicine, The Institute of Medical Science, The University of Tokyo, 4-6-1 Shirokanedai, Minato-ku, Tokyo, 108-8639 Japan, Taito Nishino, Ph.D., Planning and Development Division, Head Office, Nissan Chemical Corporation, 2-5-1 Nihonbashi, Chuo-ku, Tokyo, 103-6119 Japan. These authors contributed equally to this study (Shun Uemura, Masayuki Yamashita).

## Abstract

The distinctive milieu of the bone marrow (BM), known as the BM niche, supports hematopoietic stem cells (HSCs) and serves as a foundation for hematopoietic regeneration. Myeloablative stress disrupts not only hematopoietic stem and progenitor cells but also essential BM niche components, including endothelial cells (ECs) and mesenchymal stromal cells (MSCs); disruption of the latter impairs efficient hematopoietic recovery. However, therapeutic strategies targeting niche restoration remain largely underdeveloped. Here, we demonstrate that the Hippo pathway effectors YAP/TAZ are critical for enabling ECs and MSCs to respond to BM injury, and that YAP/TAZ activation accelerates BM niche recovery, thereby promoting hematopoietic regeneration. We found that YAP/TAZ are rapidly activated in both MSCs and ECs following myeloablative stress, maintaining MSC multipotency and orchestrating vascular remodeling. Mechanistically, YAP/TAZ function as transcriptional hubs in MSCs, regulating key transcriptional factors such as *Ebf1* and *Ebf3*. This regulation preserves MSC identity by preventing osteogenic and fibrogenic differentiation while promoting the expression of hematopoietic factors such as *Cxcl12* and angiogenic factors. In parallel, YAP/TAZ signaling in ECs facilitates pro-angiogenic responses and drives sinusoidal vessel remodeling after injury. These YAP/TAZ-mediated niche responses are essential for HSC retention and hematopoietic regeneration following diverse myelosuppressive therapies. Notably, pharmacological activation of YAP/TAZ enhances BM niche reorganization and augments hematopoietic regeneration following myeloablative therapies. These findings establish YAP/TAZ as central regulators of BM niche resilience, providing a rationale for niche-targeted therapeutic strategies to enhance hematopoietic regeneration.

**Key points:**

▪ YAP/TAZ enable BM niche cells to sense injury and restore their structure and function, thereby promoting hematopoietic regeneration.
▪ Pharmacological activation of YAP/TAZ enhances BM niche resilience and accelerates hematopoietic recovery after myelosuppressive therapies.

## Introduction

The BM is the central organ of hematopoiesis in adult mammals, where HSCs reside and generate virtually all types of blood cells throughout life^1^. Hematopoiesis is also regulated by non-hematopoietic cellular components in the BM, such as ECs, which constitute the vasculature of arterioles and sinusoids, and MSCs, which mainly exist adjacent to the vasculature and supply osteo- and adipo-lineage cells^2^. Although the BM is a highly regenerative tissue capable of regenerating blood cells after ionizing radiation (IR) and chemotherapy, such myeloablative damage causes dynamic changes in the BM niche, including dilation and leakiness of the sinusoidal vasculature^3–6^ as well as the depletion and dysfunction of MSCs^6–8^. These changes limit the efficient recovery of the hematopoietic system and curb the efficacy of HSC transplantation and chemotherapy^4,7,8^. While much is known about how BM niche cells support hematopoiesis under steady-state conditions, their role in naturally restoring the damaged BM niche remains poorly understood. Thus, therapeutic approaches that leverage niche recovery have yet to be developed.

The Hippo pathway is an evolutionarily conserved signaling cascade that acts as a cellular sensor of tissue injury^9^. It is regulated by diverse upstream inputs, including mechanical forces, extracellular matrix (ECM) stiffness, cell-cell contact, cell adhesion, and signal transduction^9^. Under steady-state conditions, these extracellular cues are relayed through multiple mediators of the Hippo pathway to activate the downstream core kinases LATS1/2, which in turn phosphorylate the central transcriptional effectors YAP and TAZ (also known as YAP1 and WWTR1, respectively), leading to their cytoplasmic retention or degradation^9,10^. Upon tissue damage, disruption of extracellular cues leads to inactivation of the Hippo pathway, resulting in YAP/TAZ activation and translocation to the nucleus and subsequent upregulation of their target genes critical for tissue regeneration^9,10^.

Compared to their well-established pro-regenerative role in solid organs, the functions of YAP/TAZ in hematopoiesis remain controversial. Earlier evidence suggests that while YAP regulates HSC development in response to the biomechanical forces from blood flow in the dorsal aorta^11^, YAP/TAZ are dispensable for both physiological and malignant hematopoiesis in adult mice^12,13^. By contrast, recent studies have implicated YAP in erythroid regeneration^14^, thrombopoiesis in immune thrombocytopenia^15^, and the control of HSC fitness following myeloablation^16^, while TAZ has been shown to protect aging HSCs from functional decline^17^. Moreover, YAP/TAZ have been shown to regulate angiogenesis^18,19^ and fibrosis^20^, and their activity increases in BM MSCs as the BM stiffness increases with age^21,22^. However, their role in restoring the BM niche following injury remains largely unexplored. Here, we uncover an unexpected requirement for YAP/TAZ in orchestrating BM niche function and regeneration. Furthermore, we demonstrate that pharmacological activation of YAP/TAZ within the niche enhances hematopoietic regeneration.

## Materials and methods

### Mice

Eight-week-old C57BL/6J mice (B6-CD45.2) were purchased from SLC Japan, and C57BL/6 mice congenic for the CD45 locus (B6-CD45.1) from Sankyo Laboratory Services. *Yap1^flox/flox^* mice were generated using *Yap1^flox/flox^* ES cells from the Knockout Mouse Project Repository (UC Davis, CA, USA), and *Taz^flox/flox^* mice were kindly provided by Dr. Jeffrey Wrana (Lunenfeld-Tanenbaum Research Institute). For generating *Yap/Taz*^ΔEC^, *Yap/Taz*^ΔMSC^, and *Yap/Taz*^ΔHC^ mice, we crossed *Yap1^fl/fl^* and *Taz^fl/fl^* mutants into *Cdh5-CreER^T2^* transgenic mice, *Ebf3-CreER^T2^* mice, and *Rosa26-CreER^T2^* mice (obtained from TaconicArtemis GmbH), respectively. To achieve EC-specific *Yap1/Taz* deletion (*Yap/Taz*^ΔEC^), 8–10-week-old *Cdh5-CreER^T2-/+^;Yap1^fl/fl^;Taz^fl/fl^*mice were injected intraperitoneally with 100 µl tamoxifen (Sigma-Aldrich) dissolved in corn oil (Sigma-Aldrich) at a concentration of 10 mg/ml for five consecutive days. *Yap/Taz*^ΔEC^ mice were analyzed 4-5 weeks after the last tamoxifen injection unless otherwise indicated. To obtain MSC-specific *Yap1/Taz* deletion (*Yap/Taz*^ΔMSC^), 8–10-week-old *Ebf3-CreER^T2-/+^; Yap1^fl/fl^;Taz^fl/fl^*mice were fed with a diet containing 400 mg/kg tamoxifen citrate (Fujifilm) (TAM food) for 14 days. Immediately after 2 weeks of TAM food administration, *Yap/Taz*^ΔMSC^ were analyzed or interventions for experiments were started. All mice were bred in the animal experiment facility at The Institute of Medical Science, The University of Tokyo (IMSUT). All experiments using mice followed the institutional guidelines for using laboratory animals and were approved by the Review Board for Animal Experiments of IMSUT (approval ID K23-46).

### Data availability

The bulk-RNA-seq, scRNA-seq, and ATAC-seq data were deposited at DNA Data Bank of Japan (DDBJ) under an accession number DRA020154.

## Results

### Niche-selective YAP/TAZ activity senses BM injury

To characterize the structural and functional changes in the BM niche upon myeloablative injury, we performed whole-mount imaging of femurs isolated from wild-type (WT) mice over the course of 14 days following sublethal IR (5 Gy). As reported previously^3,5,23^, endomucin-expressing (Emcn^+^) sinusoidal vessels of the BM exhibited drastic changes, including vascular dilation and leakiness, within 14 days before recovery (Figure 1A, 1B, Supplemental Figure 1A). Colony-forming unit-fibroblast (CFU-F) assays of whole BM cells revealed a significant reduction in CFUs following sublethal IR (Figure 1C). This, together with the previously reported marginal recovery of CFU-F activity by 4 months following lethal IR^7^, indicates incomplete recovery of MSCs in the BM following injury.

**Figure 1.**
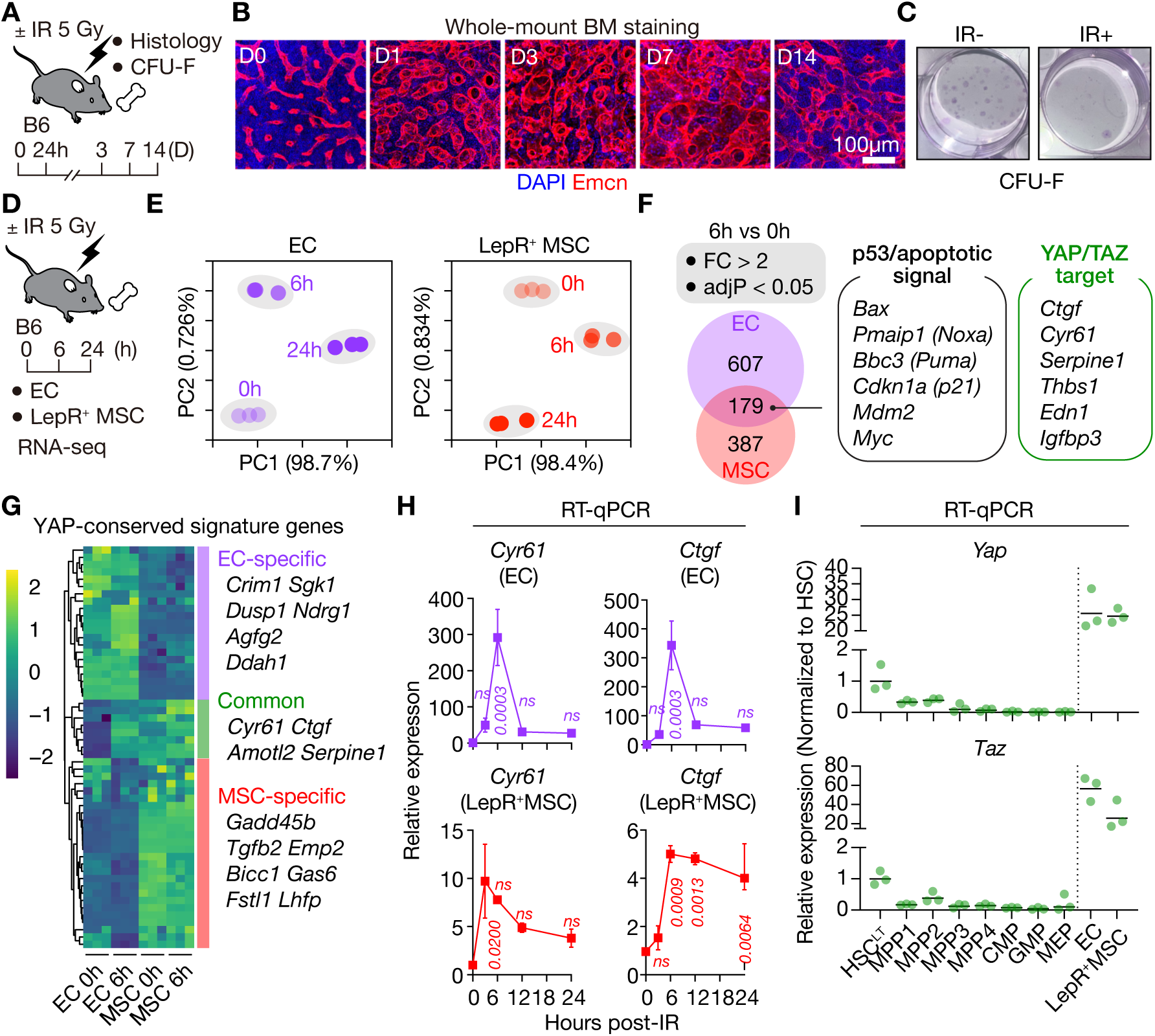
Identification of YAP/TAZ as niche-specific regulators under myeloablative stress. (A) Experimental design for BM analysis in WT mice subjected to sublethal IR (5 Gy). (B) Representative whole-mount images of femurs post-IR, stained with Endomucin (Emcn, red) and DAPI (blue). Scale bars, 100 µm. (C) Representative images of CFU-F assays performed using BM cells under steady-state conditions (IR–) and 24 hours post-IR (IR+). (D) Overview of RNA-seq analysis performed on BM ECs and LepR⁺ MSCs isolated at the indicated time points after 5 Gy IR (n = 3). (E) Principal component analysis (PCA) of transcriptomes of ECs and MSCs over time post-IR. (F) Venn diagrams showing the overlap of significantly upregulated genes (fold change (FC) > 2, false discovery rate (FDR) < 0.05) between 0- and 6-hours post-IR. (G) Heatmap showing Z-scores of normalized transcript per million (TPM) values for genes from the YAP conserved signature (MSigDB: CORDENONSI_YAP_CONSERVED_SIGNATURE). IR-responsive genes shared between ECs and MSCs are highlighted as “common”. Genes preferentially expressed in ECs and MSCs were denoted as “EC-specific” and “MSC-specific”, respectively. (H) RT-qPCR quantification of *Cyr61* and *Ctgf* transcript levels in ECs and MSCs at the indicated time points after IR, measured by RT-qPCR (n = 3). (I) RT-qPCR quantification of *Yap* and *Taz* transcript levels in BM hematopoietic and niche populations at steady state (n = 3). Data are shown as mean ± SEM. P-values were calculated using one-way ANOVA with Holm-Sidak post-hoc test (H). n.s., not significant.

To elucidate the common molecular mechanisms underlying recovery of the BM niche, we performed RNA sequencing (RNA-seq) of sinusoidal ECs and LepR⁺ MSCs isolated from the BM before and after IR (Figure 1D, Supplemental Figure 2). We focused on LepR^+^ MSCs, as they serve as the main source of HSC-supporting factors among BM stromal cells^24^. Principal component analysis (PCA) revealed time-dependent transcriptional changes in both fractions following IR (Figure 1E). Of 1,173 genes upregulated in ECs or LepR⁺ MSCs upon IR, 179 genes were commonly induced, including those associated with p53 signaling and apoptosis (Figure 1F, Supplemental Table 1). Strikingly, several canonical YAP/TAZ target genes, including *Ctgf*, *Cyr61*, and *Serpine1*, were also commonly upregulated (Figure 1F). Moreover, among the public YAP-conserved signature genes^25^, *Ctgf* and *Cyr61* were consistently upregulated in ECs and LepR⁺ MSCs post-IR (Figure 1G). Quantitative reverse transcription PCR (RT-qPCR) analysis further revealed the temporal dynamics of YAP/TAZ activity, showing that *Cyr61* and *Ctgf* were rapidly and robustly induced in both ECs and LepR⁺ MSCs after IR, peaking at approximately 6 hours, declining by 12 hours, yet remaining elevated above baseline for at least 24 hours (Figure 1H).

Since controversial results have been reported on the role of YAP/TAZ in hematopoiesis^12–17,26^, we next compared the expression of Hippo pathway genes between hematopoietic progenitors and BM niche cells. RT-qPCR analysis demonstrated that *Yap*/*Taz* are highly expressed in sinusoidal ECs and LepR⁺ MSCs, while their expression is much lower in hematopoietic cells, including HSCs (Figure 1I, Supplemental Figure 2). The upstream regulators *Lats1* and *Lats2* are abundantly expressed in ECs and LepR⁺ MSCs, with *Lats1* also being expressed in HSCs and multipotent progenitors (Supplemental Figure 1B). Reanalysis of publicly available single-cell RNA-seq (scRNA-seq) data further supported the preferential expression of Hippo pathway genes in endothelial and stromal cells over hematopoietic cells (Supplemental Figure 1C–1E). Moreover, canonical YAP/TAZ target genes—*Ctgf*, *Cyr61*, and *Amotl2*—exhibited substantially higher expression in MSCs than in ECs (Supplemental Figure 1E). In line with this, immunofluorescence imaging of *Ebf3-CreER^T2^; LSL-tdTomato* (*Ebf3-tdTomato*) femurs, where tdTomato expression marks MSCs that largely overlap with CXCL12-abundant reticular (CAR) cells and LepR⁺ cells^27^, revealed that YAP/TAZ proteins are expressed in the nuclei of steady-state and irradiated Ebf3-tdTomato^+^ MSCs (Supplemental Figure 1F, 1G), indicating that YAP/TAZ are somewhat activated in steady-state MSCs and their activity is further enhanced after injury. By contrast, YAP/TAZ protein was barely detected in sinusoidal ECs at steady state but appeared increased upon IR (Supplemental Figure 1G), consistent with the idea that YAP/TAZ are inactive in sinusoidal ECs at steady state but activated upon BM injury.

### YAP/TAZ are required in MSCs for HSC retention in the BM

The above findings indicate that YAP/TAZ not only play a homeostatic role in steady-state MSCs but also act as an early and common sensor of injury within the BM niche. To consolidate this idea by functional data, we generated a series of conditional *Yap/Taz* knockout mouse models where tamoxifen administration selectively deletes *Yap/Taz* in hematopoietic cells (*Yap/Taz*^ΔHC^), ECs (*Yap*/*Taz*^ΔEC^), and MSCs (*Yap/Taz*^ΔMSC^) (Figure 2A, 2C, 2F, Supplemental Figure 3A, 3B, 3C). Consistent with previous reports indicating a limited role for YAP/TAZ in hematopoietic cells^12,13^, *Yap/Taz*^ΔHC^ mice showed minimal changes in steady-state hematopoiesis (Figure 2B, Supplemental Figure 3D). Conversely, *Yap/Taz*^ΔEC^ mice exhibited mild cytopenia and myeloid-biased hematopoiesis (Supplemental Figure 3E), although their HSC function was largely unaffected (Figure 2D, Supplemental Figure 3F). By contrast, while no significant change was observed in the peripheral blood (PB) (Supplemental Figure 3G), immunophenotypic and functional HSC numbers were markedly reduced in the BM of *Yap/Taz*^ΔMSC^ mice, resulting in impaired repopulating capacity in competitive repopulation assays (Figure 2G, Supplemental Figure 3H). Whole-mount imaging of the BM revealed no significant alterations in the structure of sinusoidal vessels in *Yap/Taz*^ΔEC^ and *Yap/Taz*^ΔMSC^ femurs (Figure 2E, 2H). Furthermore, the spatial relationship between Ebf3-tdTomato^+^ MSCs and sinusoidal vessels remained unchanged in *Yap/Taz*^ΔMSC^ BM (Supplemental Figure 3I).

**Figure 2.**
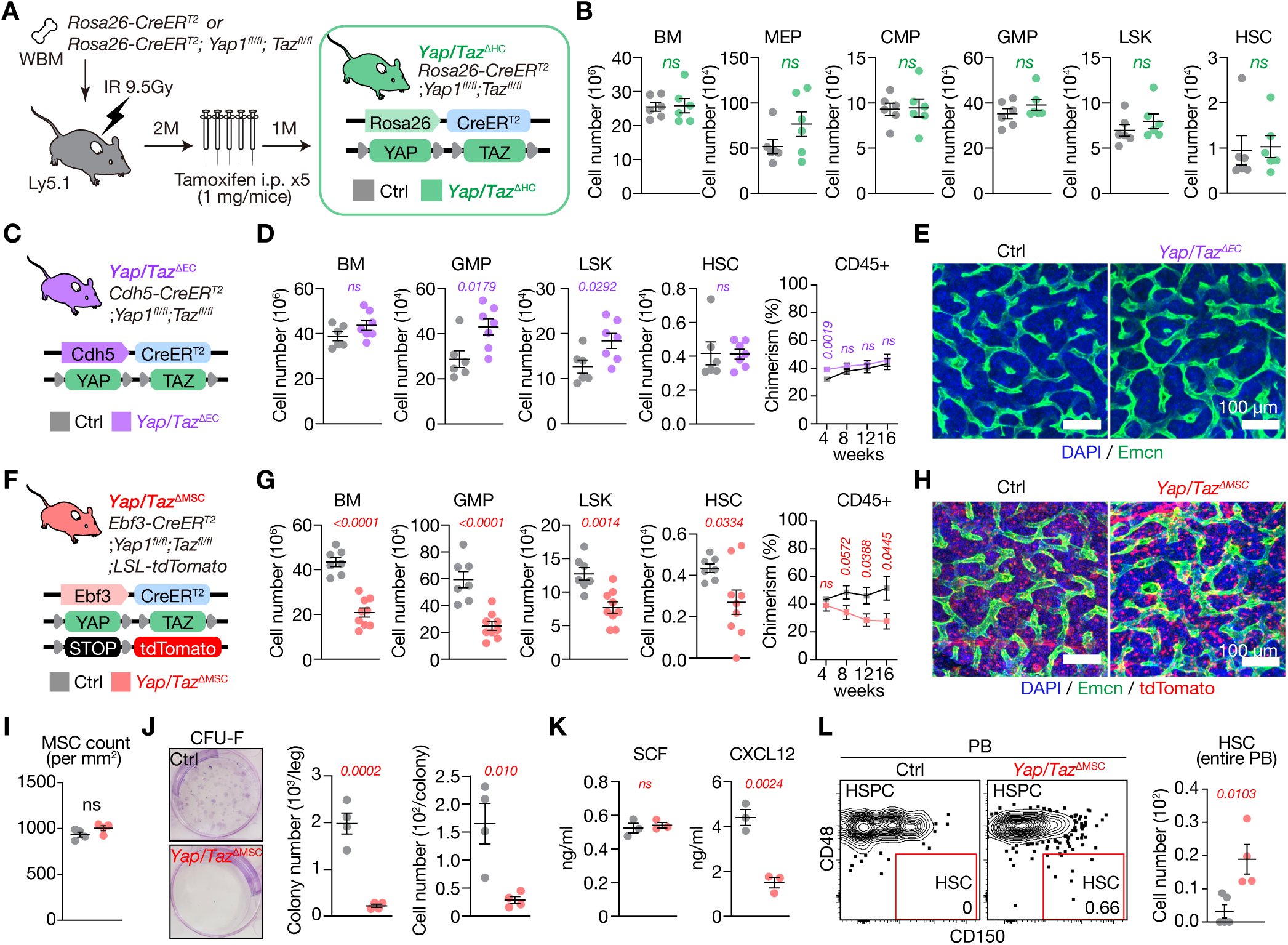
Role of YAP/TAZ in the steady-state BM niche. (A) Experimental design for conditional deletion of *Yap* and *Taz* in hematopoietic chimeras transplanted with BM cells from *Rosa26-CreER^T2^;Yap1^fl/fl^;Taz^fl/fl^*mice. (B) Absolute cell numbers of total BM, MEP, GMP, CMP, LSK, and HSC populations in a unilateral leg of *Yap/Taz*^ΔHC^ mice (n = 6). (C, F) Schematic diagrams for conditional deletion of *Yap* and *Taz* in ECs (C) and MSCs (F). (E, H) Representative whole-mount images of femurs from control (Ctrl), *Yap/Taz*^ΔEC^ (E), and *Ebf3-tdTomato^+^ Yap/Taz*^ΔMSC^ (H) mice showing signals of Emcn (green), Ebf3-tdTomato (red), and DAPI (blue). Scale bars, 100 µm. (D, G) Absolute cell numbers of total BM, GMP, LSK, and HSC populations in a unilateral leg of *Yap/Taz*^ΔEC^ (D, Ctrl, n = 5; *Yap/Taz*^ΔEC^, n = 7) and *Yap/Taz*^ΔMSC^ (G, Ctrl, n = 5; *Yap/Taz*^ΔMSC^, n = 7) mice (left). Donor chimerism (CD45.2/CD45.1+CD45.2) in PB following competitive BM transplantation of *Yap/Taz*^ΔEC^ (D, Ctrl, n = 5; *Yap/Taz*^ΔEC^, n = 8) and *Yap/Taz*^ΔMSC^ (G, Ctrl, n = 5; *Yap/Taz*^ΔMSC^, n = 8) BM cells along with competitor BM cells. (I) Quantification of Ebf3-tdTomato^+^ MSCs per unit area (/mm^2^) of BM section (n = 4). (J) Representative images of CFU-F colonies (left) and quantification of colony number and cell number per colony generated from Ctrl and *Yap/Taz*^ΔMSC^ BM (right; n = 4). (K) SCF (left) and CXCL12 (right) protein levels in BM fluid (n = 3). (L) Representative FACS plots (left) and total HSC counts in the entire PB from Ctrl (n = 5) and *Yap/Taz*^ΔMSC^ (n = 4) mice. Data are shown as mean ± SEM. P-values were calculated using two-tailed Student’s *t*-test. n.s., not significant.

As MSCs provide cues responsible for survival and retention of HSCs in the BM^28,29^, we characterized MSC identity and function following YAP/TAZ loss. Although the absolute number of MSCs was not significantly altered in *Yap/Taz*^ΔMSC^ BM, their colony-forming capacity was significantly reduced (Figure 2I, 2J). Importantly, CXCL12—a chemokine essential for HSC retention and maintenance—was markedly decreased in the BM fluid from *Yap/Taz*^ΔMSC^ mice, whereas stem cell factor (SCF), a key growth factor for HSCs, remained unchanged (Figure 2K). Consistent with this finding, circulating HSCs were significantly increased in the PB of *Yap/Taz*^ΔMSC^ mice (Figure 2L). Together, these results demonstrate that while YAP/TAZ in hematopoietic cells and ECs are largely dispensable under steady-state conditions, the basal activity of YAP/TAZ in MSCs is required for HSC retention in the BM.

### YAP/TAZ prevent MSCs from premature differentiation after injury

Next, we aimed to clarify the biological consequences of YAP/TAZ activation in the BM niche after myeloablative injury. Whole-mount BM imaging of 5 Gy-irradiated *Yap/Taz*^ΔEC^ and *Yap/Taz*^ΔMSC^ femurs revealed pronounced dilation of sinusoidal vessels in *Yap/Taz*^ΔEC^ BM, whereas it appeared attenuated in *Yap/Taz*^ΔMSC^ BM (Figure 3A–3D), suggesting that YAP/TAZ in MSCs and ECs coordinately remodel sinusoidal vessels following BM injury. While the high lethality of *Yap/Taz*^ΔEC^ mice precluded analysis beyond 10 days post-IR^30^, recovery of hematopoietic cells, including HSCs, was compromised in *Yap/Taz*^ΔMSC^ mice (Supplemental Figure 4A-4C).

**Figure 3.**
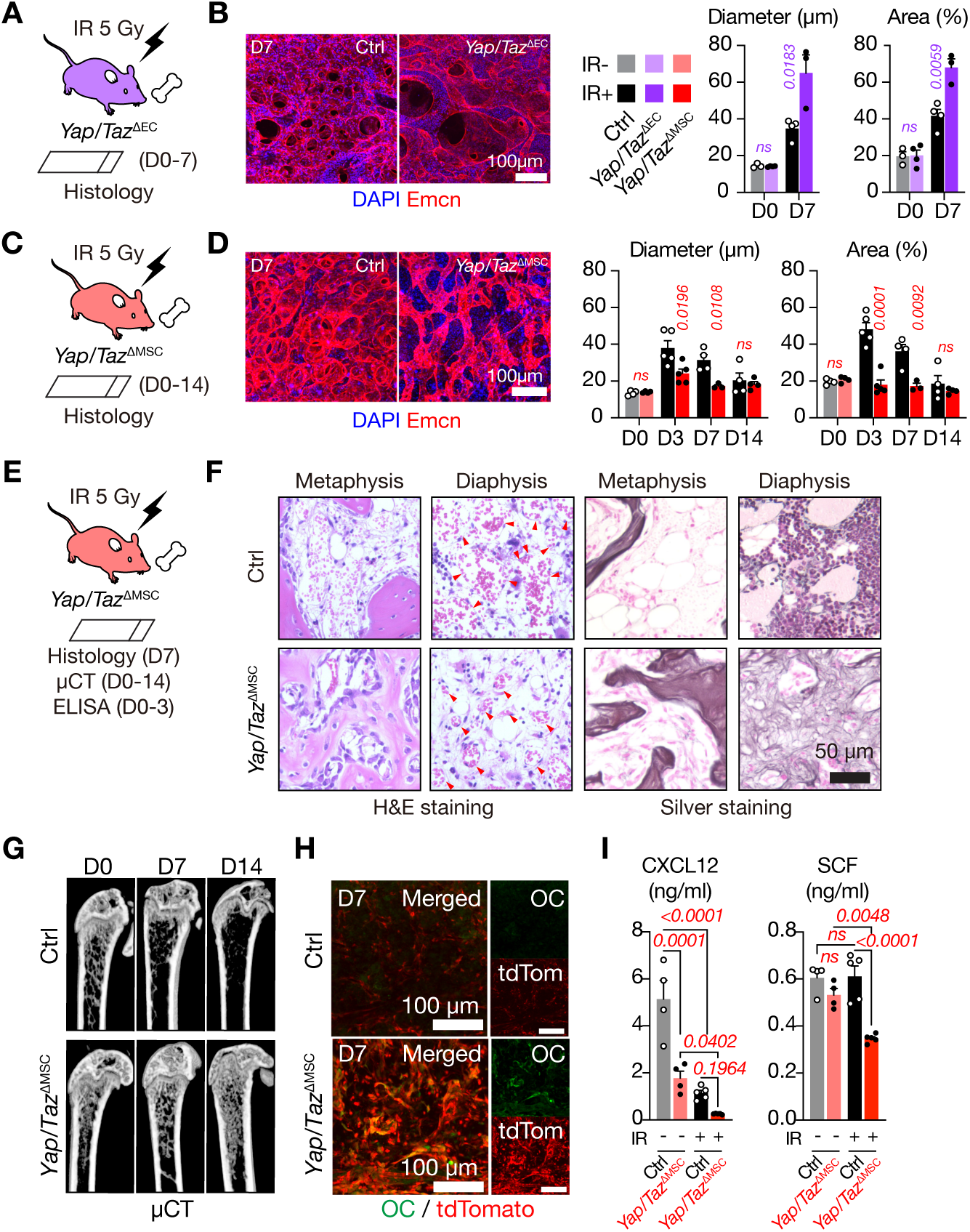
YAP/TAZ activity regulates BM niche dynamics during myeloablative stress. (A, C) Experimental design for histological analysis of BM in *Yap/Taz*^ΔEC^ (A) and *Yap/Taz*^ΔMSC^ (C) mice. (B, D) Left: Representative whole-mount images of femurs from Ctrl, *Yap/Taz*^ΔEC^ (B) and *Yap/Taz*^ΔMSC^ (D) mice 7 days after IR, showing signals of Emcn (red) and DAPI (blue). Scale bars, 100 µm. Right: Quantification of diameter and area of Emcn^+^ vessels at the indicated time points after IR (n= 3–4). (E) Experimental design for BM analysis of *Yap/Taz*^ΔMSC^ mice. (F) Representative hematoxylin and eosin (H&E) and silver staining images of Ctrl and *Yap/Taz*^ΔMSC^ femurs at day 7 post-IR. Arrowheads in H&E-staining sections of the diaphysis indicate blood vessels. (G) Representative micro-computed tomography (μCT) images of Ctrl and *Yap/Taz*^ΔMSC^ femurs. (H) Confocal images of BM sections from *Ebf3-tdTomato* Ctrl and *Yap/Taz*^ΔMSC^ femurs showing signals of Osteocalcin (OC, green) and Ebf3-tdTomato (red). Scale bars, 100 µm. (I) CXCL12 and SCF protein levels in Ctrl and *Yap/Taz*^ΔMSC^ BM fluids before and 3 days after IR. (n = 4–6). Data are shown as mean ± SEM. P-values were calculated using two-tailed Student’s *t*-test (B, D) and one-way ANOVA with Holm-Sidak post-hoc test (I). n.s., not significant.

To gain more insights into how YAP/TAZ in BM MSCs support hematopoietic recovery, we examined the histology of *Yap/Taz*^ΔMSC^ BM at day 0–14 post-IR (Figure 3E, 3F). Consistent with the immunostaining data shown in Figure 2H, steady-state *Yap/Taz*^ΔMSC^ BM exhibited no apparent histological abnormalities (Supplemental Figure 4D). However, IR-induced sinusoidal dilation was markedly attenuated in *Yap/Taz*^ΔMSC^ BM, particularly in the diaphysis (Figure 3F, Supplemental Figure 4D). Unexpectedly, this was closely associated with extensive osteosclerosis characterized by increased trabecular bone formation and fibrosis (Figure 3F, 3G, Supplemental Figure 4D). Immunostaining further revealed a substantial increase in tdTomato^+^ cells expressing osteocalcin (OC), a marker of osteoblasts, indicating enhanced osteogenic differentiation (Figure 3H, Supplemental Figure 4E). In line with this, levels of CXCL12 and SCF, the two key HSC niche factors supplied predominantly by MSCs, were significantly reduced in *Yap/Taz*^ΔMSC^ BM following IR (Figure 3I).

To comprehensively understand how YAP/TAZ regulate MSC fate, we performed scRNA-seq on BM CD45^−^ Ter119^−^ non-hematopoietic cells isolated from *Ebf3-tdTomato* control and *Yap/Taz*^ΔMSC^ mice (Figure 4A). Unsupervised clustering identified eight distinct cell clusters, including two EC clusters and six mesenchymal lineage clusters (Figure 4B). Based on established marker gene expression, two MSC clusters were annotated as adipogenic CAR cells (Adipo-CARs) and osteogenic CAR cells (Osteo-CARs) (Figure 4B, Supplemental Figure 5A). While the overall composition of mesenchymal cells remained largely unchanged under steady-state conditions, a substantial increase in the proportion of Osteo-CARs, osteoblasts, and fibroblast-like (fibrotic) cells was observed in *Yap/Taz*^ΔMSC^ BM by day 3 post-IR (Supplemental Figure 5B). Further analysis on *Ebf3-tdTomato*^+^ cells revealed a similar lineage shift (Figure 4C, Supplemental Figure 5C), confirming that *Yap/Taz*-deficient MSCs undergo accelerated differentiation toward osteoblastic and fibrotic cells following BM injury. Moreover, *Cxcl12* and *Scf* (also known as *Kitl*) were significantly downregulated in *Yap/Taz*^ΔMSC^ MSCs (Figure 4D). Collectively, these findings indicate that MSCs rely on YAP/TAZ to maintain their identity and to produce HSC-supporting factors, particularly following BM injury.

**Figure 4.**
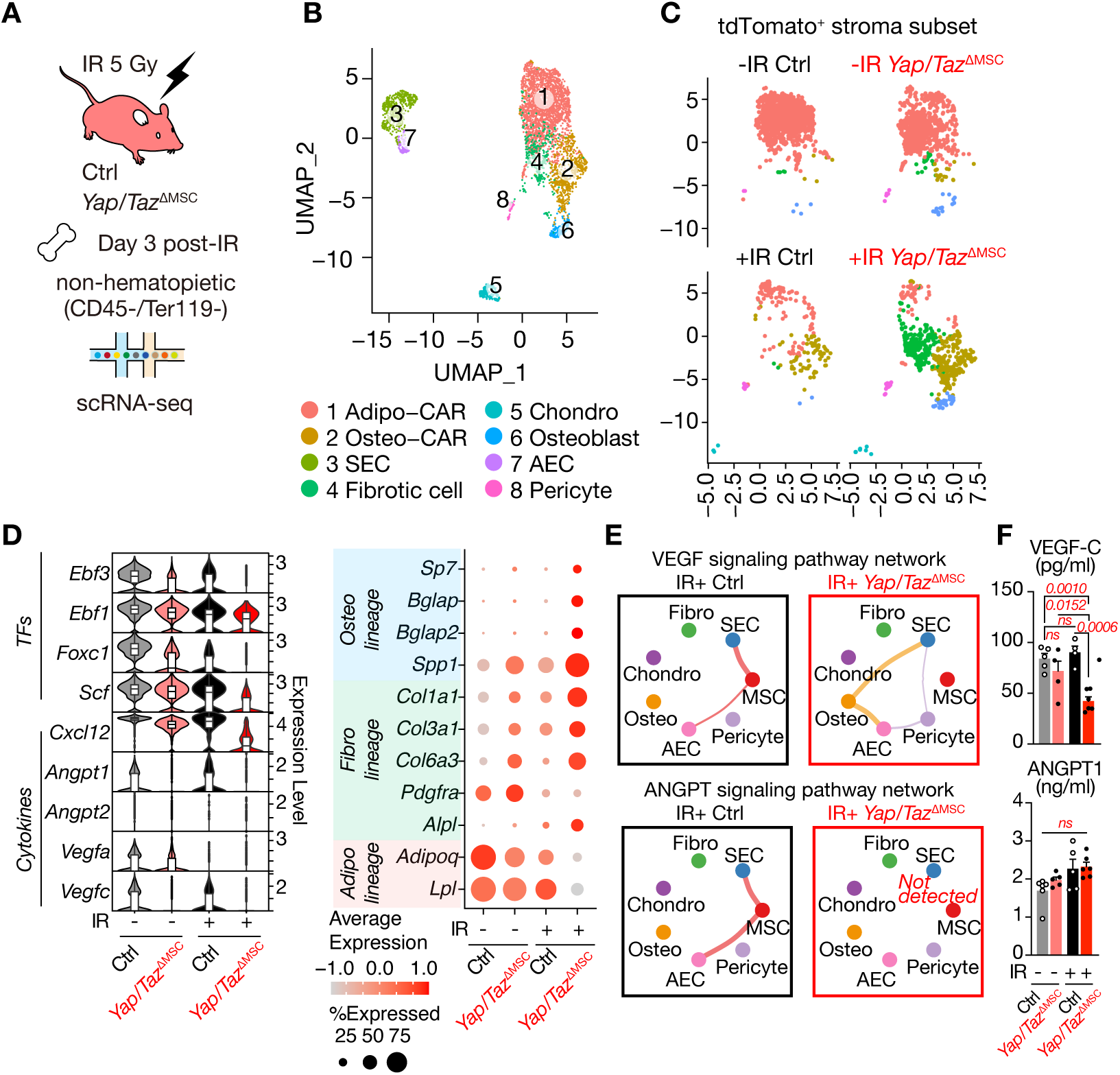
YAP/TAZ are key regulators of MSC differentiation and function. (A) Experimental design for scRNA-seq analysis of non-hematopoietic cells isolated from Ctrl and *Yap/Taz*^ΔMSC^ BM. (B) UMAP projection of eight cell clusters of BM niche cells, derived from merged scRNA-seq data (4,106 cells) from the four samples shown in (D. C) UMAP plots showing eight cell clusters of *Ebf3-tdTomato^+^* MSCs in Ctrl and *Yap/Taz*^ΔMSC^ BM before and 3 days after IR. (D) Violin and dot plots showing the expression levels of key transcription factors, niche factors, and osteo/adipo/fibro-lineage marker genes in *Ebf3-tdTomato^+^* MSCs from Ctrl and *Yap/Taz*^ΔMSC^ mice before and 3 days after IR. (E) Circle plots of ligand-receptor interactions between BM niche populations, inferred from scRNA-seq data using CellChat. (F) VEGF-C and ANGPT1 protein levels in Ctrl and *Yap/Taz*^ΔMSC^ BM fluids before and 3 days after IR. (n = 4–6). Data are shown as mean ± SEM. P-values were calculated using one-way ANOVA with Holm-Sidak post-hoc test

### YAP/TAZ in the BM niche coordinate vascular remodeling

The restricted dilation of sinusoidal vessels in *Yap/Taz*^ΔMSC^ BM may be attributable to an increased extravascular burden, such as that imposed by hematopoietic cells and ECM components. However, BM cell numbers were comparably reduced in both control and *Yap/Taz*^ΔMSC^ mice following IR (Supplemental Figure 4B). While the osteosclerotic and fibrotic changes in the extravascular space may contribute to the restricted dilation, an alternative possibility is impaired regulation of ECs due to reduced production of angiogenic factors by MSCs, such as VEGF-C^31^ and ANGPT1^32,33^. Indeed, the expression of *Vegfc* and *Angpt1* was downregulated in *Yap/Taz*^ΔMSC^ MSCs both at steady state and after IR (Figure 4D, Supplemental Figure 5D). We then performed ligand-receptor analysis on the scRNA-seq data using the “CellChat” package to explore intercellular communication among different niche cell types (Supplemental Figure 6A). In control BM, we observed strong projections of VEGF and ANGPT signaling from MSCs to ECs. Notably, deletion of *Yap/Taz* in MSCs diminished MSC-to-EC communication via VEGF and ANGPT signaling pathways (Figure 4E, Supplemental Figure 6B). Consistent with these altered signaling interactions, protein levels of VEGF-C, but not ANGPT1, were significantly reduced in the BM fluid of irradiated *Yap/Taz*^ΔMSC^ mice (Figure 4F). Given that ANGPT1 is primarily secreted by perivascular MSCs and likely acts locally^33^, its effects may be spatially restricted and therefore not fully captured by its levels in the BM fluid.

YAP/TAZ in ECs have been shown to mediate VEGF-induced angiogenesis and vascular development^34,35^. Indeed, bulk RNA-seq on *Yap/Taz*^ΔEC^ ECs revealed downregulation of genes involved in vascular development, cell adhesion, and ECM organization under steady-state conditions (Supplemental Figure 7A–7C, Supplemental Table 2, 3). Gene set enrichment analysis (GSEA) further demonstrated that these genes were readily induced in control ECs following IR, an effect that was largely abolished in the absence of YAP/TAZ (Supplemental Figure 7D, 7E). In particular, *Apelin* (*Apln*) was upregulated in ECs upon IR in a YAP/TAZ-dependent manner (Supplemental Figure 7B, Supplemental Table 2), supporting the notion that YAP/TAZ may promote vascular regeneration through expansion of the *Apln^+^*EC population^5^. Collectively, these results suggest that YAP/TAZ are critical to engage a pro-angiogenic response in sinusoidal ECs, potentially by regulating angiogenic signals derived from MSCs.

### YAP/TAZ activate a core transcriptional network that governs MSC identity

YAP/TAZ regulate target gene expression via interaction with transcription factors, such as TEAD^9^. To gain mechanistic insight into how YAP/TAZ regulates MSC identity, we performed ATAC-seq analysis on control and *Yap/Taz*^ΔMSC^. Ebf3-tdTomato^+^ MSCs revealed reduced chromatin accessibility at genomic regions containing TEAD-binding motifs in steady-state *Yap/Taz*^ΔMSC^ MSCs (Figure 5A, 5B, Supplemental Figure 8A, Supplemental Table 4, 5). Accessibility of adipogenesis-related EBF^27,36^-binding motifs was also significantly decreased, whereas that of osteogenesis-related RUNX-binding motifs was increased in *Yap/Taz*^ΔMSC^ MSCs (Figure 5B, Supplemental Figure 8B, Supplemental Table 5). Digital footprint analysis further demonstrated reduced occupancy of TEAD at its cognate motifs (Figure 5C, 5D, Supplemental Table 6) and decreased occupancy of transcription factors implicated in adipogenic (CEBPA^37^, EBF1/3^27,36^) and osteogenic (FOXC1^38^) differentiation at their respective binding motifs in *Yap/Taz*^ΔMSC^ MSCs (Figure 5C, 5D, Supplemental Figure 8B, 8C, Supplemental Table 6). TEAD binding was identified at the promoter or upstream regulatory regions of *Ebf1/3*, *Foxc1*, *Cxcl12*, *Vegfc*, and *Angptl1* (Figure 5E, Supplemental Table 6), in line with their decreased expression in steady-state and irradiated *Yap/Taz*^ΔMSC^ MSCs (Figure 4D, Supplemental Figure 5D). Among key transcription factors, EBF1/3 appeared to be critical downstream regulators of YAP/TAZ, as reanalysis of *Lepr-Cre;Ebf1^fl/+^;Ebf3^fl/fl^* BM revealed not only progressive osteosclerosis^27^ but also fibrosis similar to that observed in *Yap/Taz*^ΔMSC^ BM (Figure 5F). Collectively, these findings underscore the central role of the YAP/TAZ–TEAD axis in MSCs in orchestrating transcriptional networks critical for maintaining MSC identity, as well as their HSC-supportive and pro-angiogenic functions (Figure 5G).

**Figure 5.**
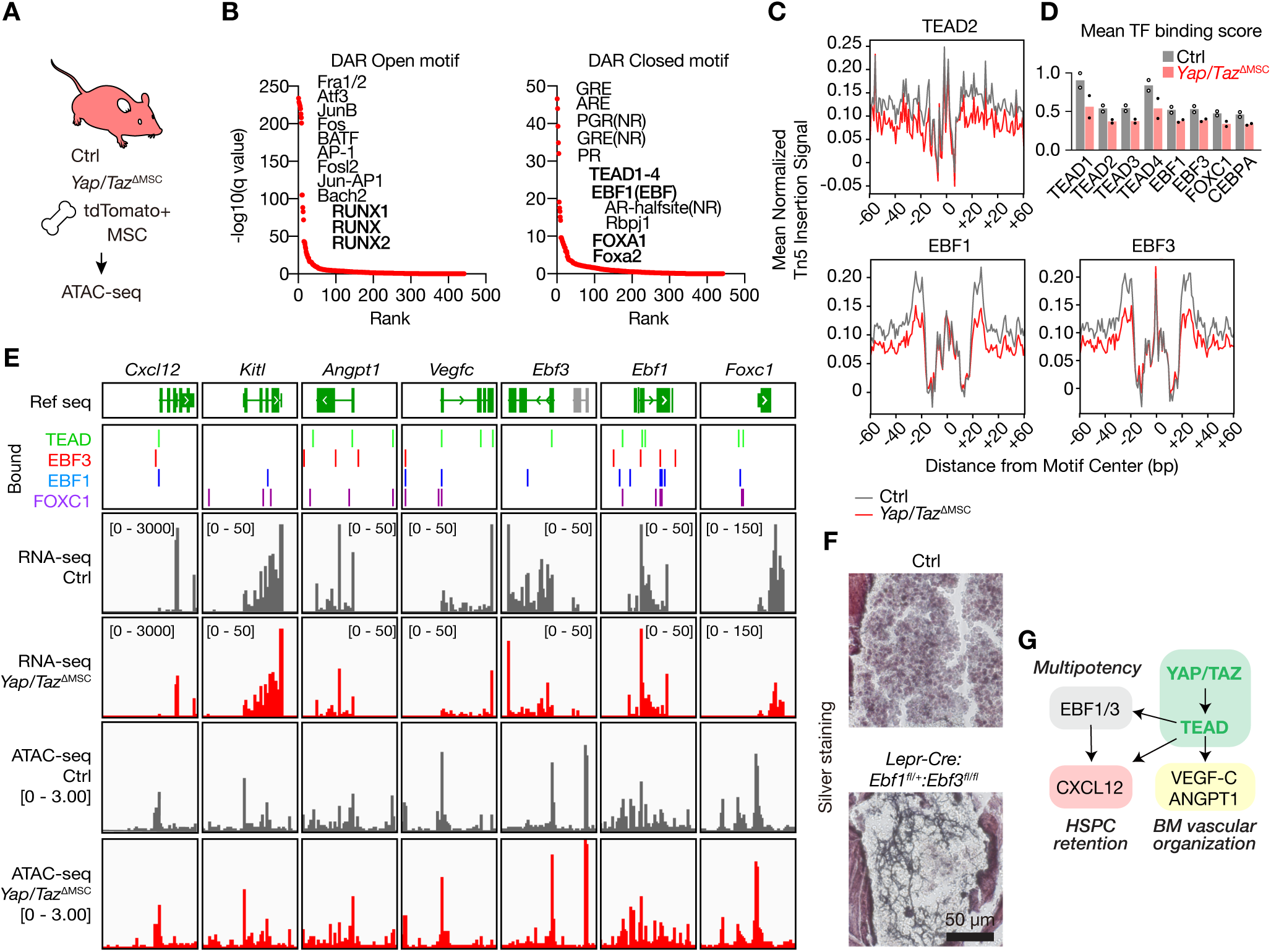
YAP/TAZ function as transcriptional network hubs in MSCs. (A) Experimental design for ATAC-seq analysis of Ebf3-tdTomato^+^ BM MSCs from Ctrl and *Yap/Taz*^ΔMSC^ mice. (B) Enrichment ranking of transcription factor binding motifs within DARs that were either opened or closed in *Yap/Taz*^ΔMSC^ MSCs. (C) Mean normalized Tn5 insertion signals around TEAD2, EBF1, and EBF3 motifs in Ebf3-tdTomato^+^ Ctrl and *Yap/Taz*^ΔMSC^ MSCs, indicating transcription factor occupancy as computed by TOBIAS. (D) Mean transcription factor binding scores (TFBS) for selected transcription factor binding motifs within consensus ATAC-seq peaks. (E) Integrative Genomics Viewer (IGV) tracks showing pseudo-bulk scRNA-seq and ATAC signals around *Cxcl12*, *Ebf1*, *Ebf3*, *Vegfc*, and *Angpt1* loci in Ctrl and *Yap/Taz*^ΔMSC^ MSCs. Gene structures are shown at the top. Positive transcription factor binding inferred from footprinting in Ctrl Ebf3-tdTomato*^+^* MSCs is highlighted as “bound”. (F) Representative silver staining images of femur sections from 34-week-old Ctrl and *Lepr-Cre;Ebf1^fl/+^;Ebf3^fl/fl^*mice. (G) Graphical summary of transcriptional regulation of MSC identity and function by the YAP/TAZ-TEAD axis.

### A novel LATS1/2 inhibitor promotes hematopoietic regeneration by enhancing niche resilience

Given the critical role of YAP/TAZ in maintaining the BM niche functionality, we hypothesized that enhancing YAP/TAZ activity could promote niche recovery and thereby facilitate hematopoietic regeneration following myeloablative injury. We recently identified GA-017, a potent small-molecule inhibitor of LATS1/2 that activates YAP/TAZ (Supplemental Figure 9A)^39^. GA-017 preferentially targets AGC family kinases, with LATS1/2 being most efficiently inhibited, as evidenced by its low half-maximal inhibitory concentration (IC_50_) values (4.10 ± 0.79 for LATS1 and 3.92± 0.42 nM for LATS2)^39^. To test our hypothesis, we developed a series of GA-017 derivatives with a similar structure and selected GA-003 based on its fourfold greater potency than GA-017 (Supplemental Figure 9A, 9B). We confirmed that GA-003 treatment enhanced nuclear translocation of YAP/TAZ in both human umbilical vein endothelial cells (HUVECs) and mouse OP-9 stromal cells (Supplemental Figure 9C). Consistent with the low expression levels of *Yap*/*Taz* in early hematopoietic progenitors (Figure 1I), GA-003 did not affect LSK cell growth *in vitro* (Supplemental Figure 9D).

Next, we administered GA-003 intraperitoneally at a dose of 1.25 mg per mouse and evaluated its effect *in vivo* (Supplemental Figure 9E). Within 6 hours post-injection, we observed significant upregulation of *Cyr61* and *Ctgf* in both BM ECs and MSCs (Supplemental Figure 9F), indicating that YAP/TAZ are activated in the BM niche. To evaluate the effect of YAP/TAZ activation on BM niche recovery, mice were exposed to 5 Gy IR and treated with eight daily doses of GA-003, beginning 30 min before IR (Figure 6A). While IR-induced myelosuppression was indistinguishable between vehicle- and GA-003-treated mice, hematopoietic recovery was significantly accelerated in GA-003-treated mice (Figure 6B, 6C, Supplemental Figure 9G). GA-017 also showed a similar effect on hematopoietic recovery, but to a lesser extent than GA-003, consistent with its lower potency for LATS1/2 inhibition (Supplemental Figure 9H–9J). Thus, the novel LATS1/2 inhibitor GA-003 potently promotes hematopoietic regeneration, likely via boosting YAP/TAZ activity in the BM niche during recovery.

**Figure 6.**
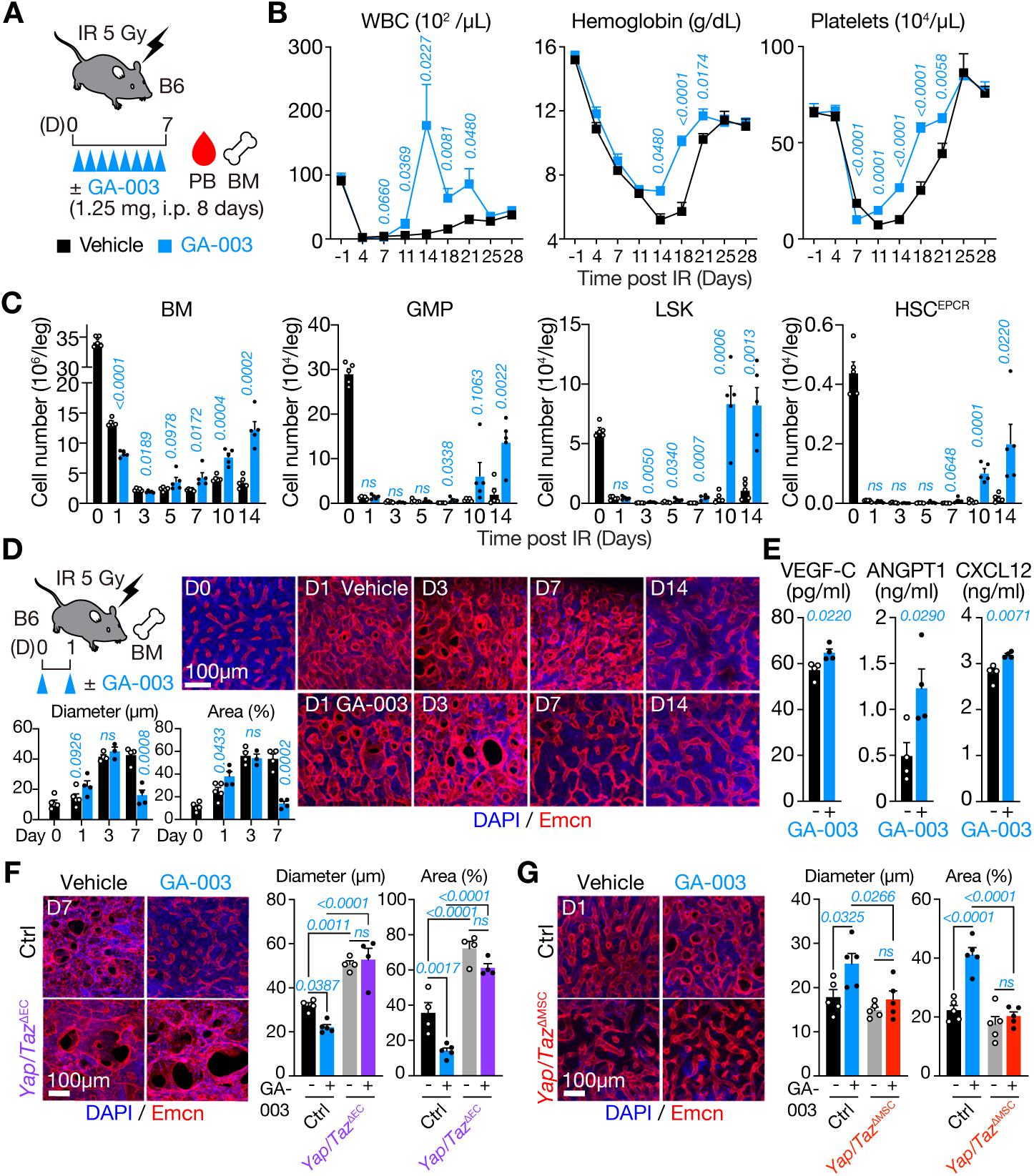
A novel LATS1/2 inhibitor enhances niche recovery and hematopoietic regeneration. (A) Experimental design for GA-003 treatment of sublethally irradiated WT mice. B, PB counts (n = 8) and (C) absolute cell numbers of total BM, GMP, LSK, and HSC^EPCR^ populations (n = 5 each) at the indicated time points post-IR. (D) Effect of GA-003 on sinusoidal vessel remodeling after IR. Representative whole-mount images of Emcn^+^ sinusoidal vessels in femurs on day 14 post-IR, showing DAPI (blue) and Emcn (red) signals. Quantification of the diameter (left) and area (right) of the Emcn^+^ sinusoidal vessels are also shown (n = 3–5). (E) ANGPT1, VEGF-C, and CXCL12 protein levels in BM fluid of Ctrl and GA-003-treated mice on day 1 post-IR (n = 4). (F-I) Effects of GA-003 on vascular remodeling in *Yap/Taz*^ΔEC^ (F) and *Yap/Taz*^ΔMSC^ (G) mice. Representative whole-mount BM images and quantification of Emcn^+^ sinusoidal vessel diameter and area in femurs on day 7 post-IR. Data are shown as mean ± SEM. P-values were calculated using two-tailed *t*-test (B, C, D, E) and one-way ANOVA with Holm-Sidak post-hoc test (F,G). n.s., not significant.

Our dose de-escalation studies revealed that the initial two doses of GA-003 were sufficient to accelerate hematopoietic recovery (Supplemental Figure 10A–10C), consistent with the notion that the BM niche relies on early YAP/TAZ activation following IR to restore its HSC-supporting function. To identify the cellular targets of GA-003, *Yap/Taz*^ΔHC^, *Yap/Taz*^ΔEC^, and *Yap/Taz*^ΔMSC^ mice were subjected to sublethal IR and treated with GA-003 (Supplemental Figure 10D, 10G, 10J). Notably, while GA-003 accelerated hematopoietic recovery in both control and *Yap/Taz*^ΔHC^ mice to a similar extent, its effects were partially diminished in *Yap/Taz*^ΔEC^ and almost completely abolished in *Yap/Taz*^ΔMSC^ mice (Supplemental Figure 10D–10L). These results support the notion that GA-003 promotes hematopoietic recovery primarily through its actions on the BM niche, particularly *Ebf3^+^* MSCs. Indeed, GA-003 treatment accelerated the entire process of BM vascular remodeling (Figure 6D), which appeared to be partly mediated by increased secretion of VEGF-C and ANGPT1 by MSCs (Figure 6E). Consistently, GA-003 failed to rescue defects in vascular remodeling not only in *Yap/Taz*^ΔEC^ mice but also in *Yap/Taz*^ΔMSC^ mice (Figure 6F, 6G). Furthermore, GA-003 enhanced VE-cadherin expression on ECs and reduced vascular leakiness, thereby promoting recovery of vascular integrity (Supplemental Figure 11A–11C).

To gain comprehensive insight into the mechanism of action of GA-003, we performed scRNA-seq on CD45^−^ Ter119^−^ non-hematopoietic BM cells isolated from GA-003-treated *Ebf3-tdTomato* mice at days 0 and 3 post-IR (Supplemental Figure 11D, 11E). In control BM, the frequency of Adipo-CAR cells declined, whereas Osteo-CAR cells increased following IR (Supplemental Figure 11F–11H). Although GA-003 had minimal impact on these cell frequencies, it moderately upregulated *Ebf3* and *Ebf1* in *Ebf3-tdTomato^+^* stromal cells after IR (Supplemental Figure 11I). Consistently, transcripts and protein levels of angiogenic factors, including VEGF-C and ANGPT1, were elevated in the BM of GA-003-treated mice (Figure 6E, Supplemental Figure 11I). In addition, GA-003 increased the expression of several adhesion molecules and ECM-related genes in ECs (Supplemental Figure 11I). Together, these results establish that pharmacological boosting of YAP/TAZ activity promotes hematopoietic regeneration by enhancing the recovery of the BM niche.

### LATS1/2 inhibition ameliorates diverse myelosuppressive conditions

In clinical settings, patients frequently experience severe myelosuppression during chemotherapy and HSC transplantation. To evaluate the therapeutic potential of LATS inhibition under such conditions, we examined the effects of GA-003 in mouse models of chemotherapy and HSC transplantation. Consistent with its effect following IR, GA-003 treatment significantly enhanced hematopoietic recovery after administration of 5-fluorouracil (5-FU), a clinically relevant myelosuppressive chemotherapeutic agent (Figure 7A). GA-003 also acted synergistically with granulocyte-colony stimulating factor (G-CSF), a cytokine widely used to treat neutropenia, to promote PB leukocyte recovery following IR-induced myelosuppression (Figure 7B). Moreover, GA-003 promoted the engraftment of both mouse (Figure 7C) and human HSCs (Figure 7D) following transplantation. Collectively, these findings demonstrate that GA-003 promotes hematopoietic regeneration across multiple myeloablative conditions, highlighting its therapeutic potential in the context of chemotherapy and HSC transplantation.

**Figure 7.**
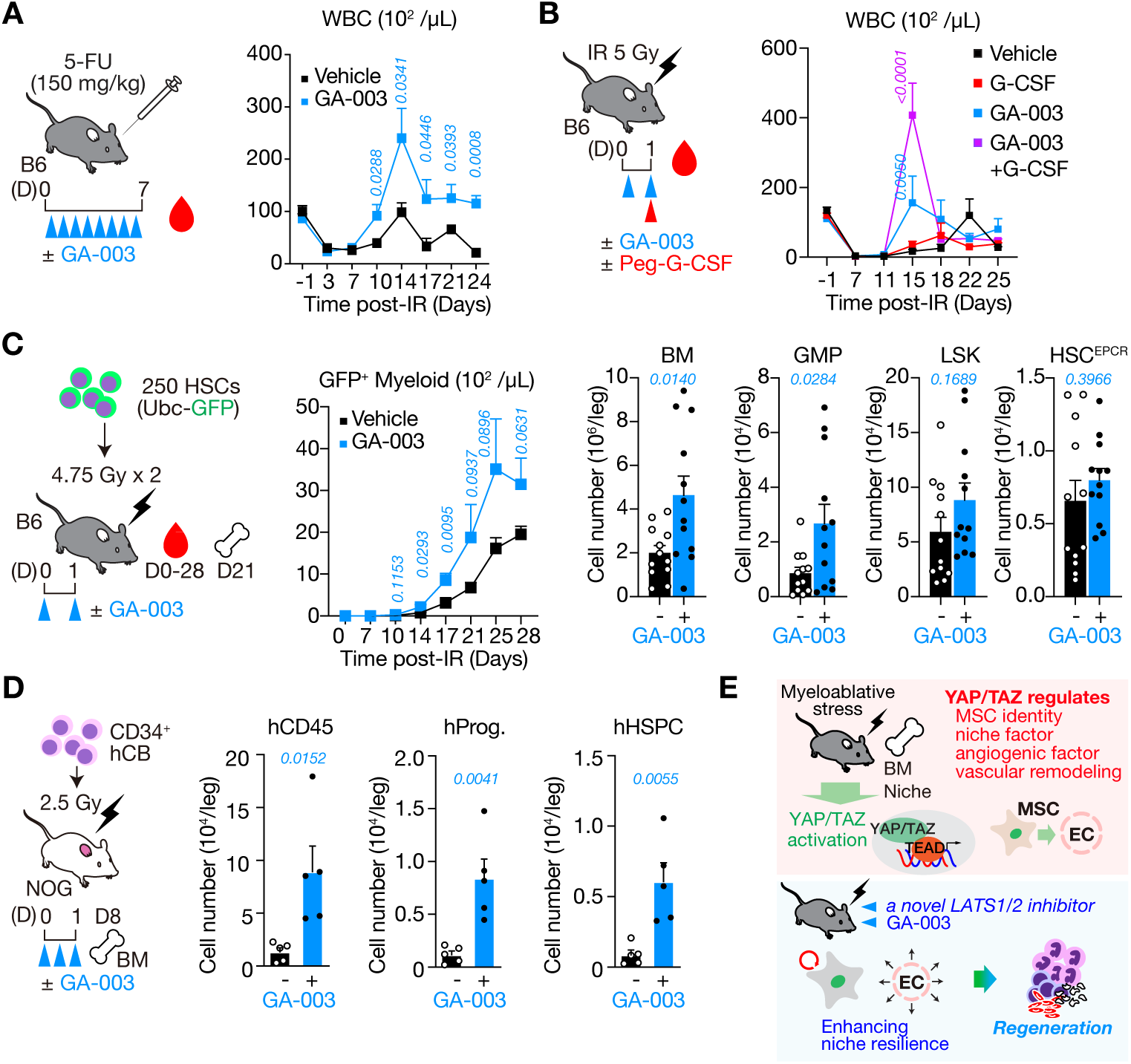
A novel LATS1/2 inhibitor accelerates recovery from various types of myelosuppression. (A) Effect of GA-003 on WBC recovery after 5-FU treatment (n = 4–5). (B) Combinatory effect of GA-003 and G-CSF on WBC recovery after IR (n = 5). (C) Impact of GA-003 injection on the mouse HSC transplantation model. GFP-expressing HSCs were transplanted into lethally irradiated mice (left). Absolute cell numbers of GFP^+^ donor-derived myeloid cells in PB (n = 5) (middle), total BM, GMP, LSK, and HSC^EPCR^ populations on day 21 post-HSC transplantation (n = 12) (right). (D) Effect of GA-003 on human HSPC repopulation in a xenograft model. Absolute numbers of human CD45^+^ hematopoietic cells, CD34^+^CD38^+^ hematopoietic progenitor cells (Prog.), and CD34^+^CD38^−^ hematopoietic stem and progenitors (HSPCs) in the BM (n = 5). (E) Graphical summary of YAP/TAZ functions and GA-003 action on the BM niche and hematopoietic regeneration. Data are shown as mean ± SEM. P-values were calculated using two-tailed Student’s *t*-test. n.s., not significant.

## Discussion

Here, we provide direct evidence that YAP/TAZ in MSCs and sinusoidal ECs, but not in hematopoietic cells, support hematopoietic regeneration via coordinated regulation of BM niche recovery after myeloablative injury (Figure 7E). Our data demonstrate that YAP/TAZ activation enables both MSCs and ECs to respond to BM injury, thereby promoting BM niche recovery. Despite their basal activity at steady state, YAP/TAZ in MSCs are further activated upon injury to preserve MSC identity and maintain their HSC-supportive and pro-angiogenic functions. By contrast, YAP/TAZ in ECs are strictly suppressed under steady-state conditions but become readily activated upon injury to drive sinusoidal vessel remodeling. Furthermore, our newly identified LATS1/2 inhibitor, GA-003, enhances BM niche resilience and promotes hematopoietic regeneration in a YAP/TAZ-dependent manner. Together, our results establish YAP/TAZ as a master regulator of BM niche resilience that can be pharmacologically targeted to promote hematopoietic regeneration after myelosuppressive therapy.

Our results identify YAP/TAZ–TEAD as a key transcriptional regulator that protects BM MSCs from premature differentiation. While previous studies have underscored the importance of various transcriptional and epigenetic regulators in maintaining MSC identity, the upstream regulatory mechanisms controlling these factors have remained largely unclear. Our data indicate that YAP/TAZ act upstream of multipotency regulators such as *Foxc1 and Ebf1/3*, as well as other key factors critical for vascular remodeling and hematopoietic regeneration (Figure 5G). FOXC1 and EBF1/3 are known to preserve CAR cell identity by repressing adipogenic and osteogenic differentiation, respectively. Deletion of *Foxc1* and *Ebf1/3* in LepR^+^ MSCs results in severe BM adipogenesis and osteosclerosis, respectively^27,38,40^. Both factors are also essential for maintaining high *Cxcl12* expression in CAR cells^27,38,41^. Our results are in line with these observations and suggest that YAP/TAZ–TEAD maintains MSC identity by directly inducing these key transcriptional regulators.

Our findings provide a molecular basis for how the two phases of sinusoidal vessel remodeling—dilation and recovery — are regulated by MSCs and ECs, respectively. YAP/TAZ in MSCs appear to regulate early sinusoidal dilation, at least partly, through the secretion of angiogenic factors involved in vascular remodeling during hematopoietic regeneration^32,33,42^. On the other hand, although post-IR lethality in *Yap/Taz*^ΔEC^ mice precluded evaluation of whether their sinusoids eventually undergo vascular recovery, the negligible effect of YAP/TAZ deficiency in ECs on sustained sinusoidal dilation suggests that they play a role in sinusoidal recovery rather than dilation. The precise mechanisms underlying this YAP/TAZ-dependent coordination between MSCs and ECs in regulating sinusoidal vessel remodeling warrant further investigation.

Given the multifaceted functions of YAP/TAZ in BM niche resilience and hematopoiesis, as well as the evolutionarily conserved nature of Hippo–YAP/TAZ signaling, we developed a niche-targeted therapeutic approach through pharmacological activation of YAP/TAZ. Although previous efforts to develop LATS1/2 inhibitors faced challenges due to broad on- and off-target side effects across organs and potential tumorigenicity^43–45^, our strategy using two doses of the reversible LATS1/2 inhibitor GA-003 successfully induced transient YAP/TAZ activation in the BM niche. This promoted niche recovery and hematopoietic regeneration following various myelosuppressive therapies in preclinical mouse models. GA-003 also showed promising effects on human cord blood HSC engraftment in immunodeficient mice; however, its impact on the human BM niche remains to be evaluated. Future studies using human BM organoids and/or non-human primate models will pave the way to advance this niche-modulating strategy toward clinical application.

## AUTHOR CONTRIBUTIONS

S.U., M.Y., T.Y-N., and A.I. conceived the project and designed the experiments. S.U., M.Y., and T.Y-N. performed experiments, analyzed data, and actively wrote the paper. A.A., T.I., S.K., Y.N.-T., M.O., K.K., and T.N. designed and performed experiments. B.R., and A.K. performed sequencing. M.N., A.S., and A.S. generated mice. T.N., K.K., and T.N. discussed the results. A.I. guided and supervised the project, secured funding, and wrote the paper. All authors edited and approved the paper.

## ACKNOWLEDGMENTS

We thank J. Wrana (Lunenfeld-Tanenbaum Research Institute) and Y. Kubota (Keio University) for *Taz^flox/flox^* and *Cdh5-CreER^T2^*mice, respectively. We are also grateful to Dr. Sandra Capellera Garcia (St. Jude Children’s Research Hospital) for her critical reading and editing of the manuscript. The super-computing resource was provided by The Human Genome Center, The Institute of Medical Science, The University of Tokyo (IMSUT). This work was supported in part by Grants-in-Aid for Scientific Research (19H05653, 24H00066, 19H05746 to AI, 23K19622, 24K19218, 24KJ0053 to SU, 22H03101 to MY) from the Japan Society for the Promotion of Science (JSPS), 21zf0127003h0001 and 243fa627001 (AI) from the Japan Agency for Medical Research and Development (AMED), and Nissan Chemical Corporation.

## DECLARATION OF INTERESTS

TN, KK, AA, and TI are employees of Nissan Chemical Corporation.

