## SupplementalFigure for "Niche-targeted therapy via YAP/TAZ activation enhances hematopoietic regeneration"

### Supplemental Figure 1

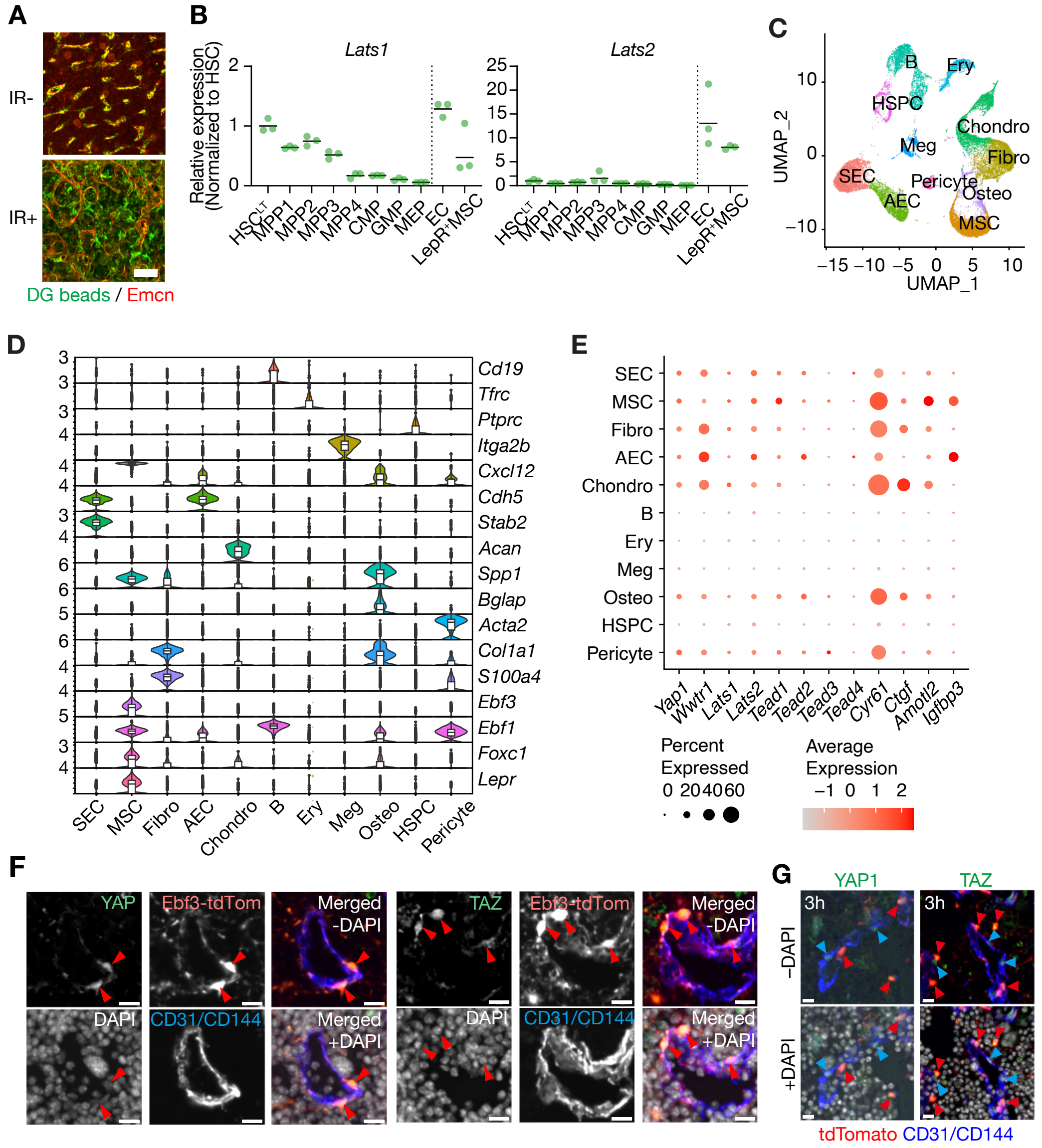


**Supplemental Figure 1 | Expression of *Yap/Taz* and their related genes in BM niche cells.** (A) Representative immunofluorescence images of femurs from mice under steady-state conditions (IR–) and on day 7 post-IR (IR+). Dragon Green (DG) beads (green), and Emcn (red). (B) RT-qPCR quantification of *Lats1*, and *Lats2* transcript levels in BM hematopoietic and niche populations (n = 3). (C) UMAP illustration of BM single-cell transcriptomes from Baryawno *et al*. (D) Stacked violin plots showing the expression of signature genes defining hematopoietic and non-hematopoietic BM cell types. (E) Dot plots showing the expression levels of Hippo pathway genes and YAP/TAZ target genes in each BM hematopoietic and non-hematopoietic component. (F) Representative confocal immunofluorescence images of *Ebf3-tdTomato* femur sections at steady state showing signals for YAP and TAZ (green), Ebf3-tdTomato (red), CD31/CD144 (blue), and DAPI (gray). Red arrowheads indicate the nuclei positive for Ebf3-tdTomato and YAP or TAZ. Scale bars, 20 μm. (G) Representative confocal immunofluorescence images of *Ebf3-tdTomato* femur sections at 3 hours post-IR, showing signals for YAP and TAZ (green), Ebf3-tdTomato (red), CD31/CD144 (blue), and DAPI (gray). Red arrowheads indicate the nuclei positive for Ebf3-tdTomato and YAP or TAZ, while blue arrowheads indicate the nuclei positive for CD31/CD144 and YAP or TAZ. Scale bars, 20 μm.

### Supplemental Figure 2

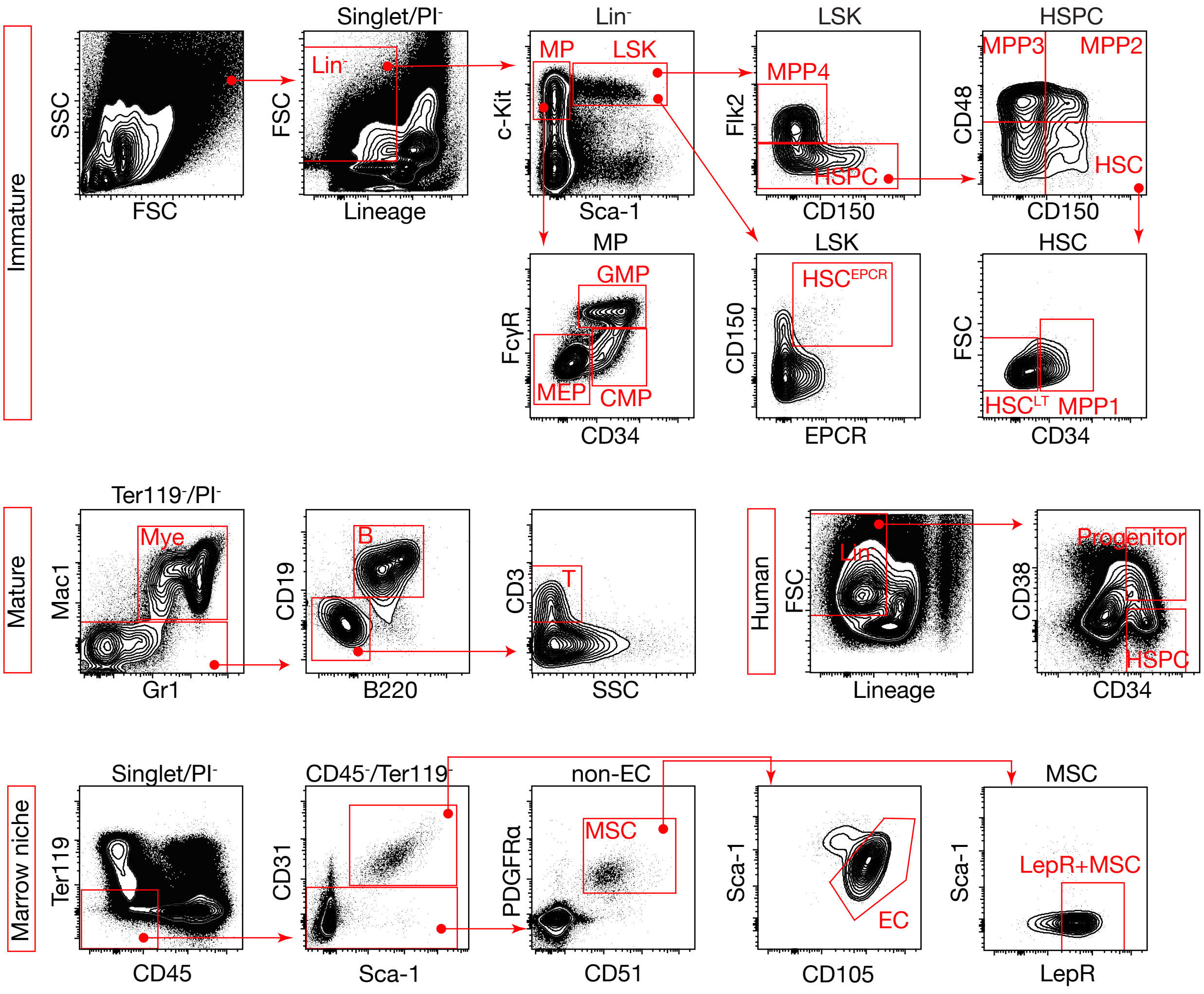


**Supplemental Figure 2 | Flow cytometric profiles of HSPCs.**
Flow cytometry profiles and gating of murine HSPCs, mature hematopoietic cells, human cord blood HSPCs, murine BM ECs, and MSCs are shown.

### Supplemental Figure 3

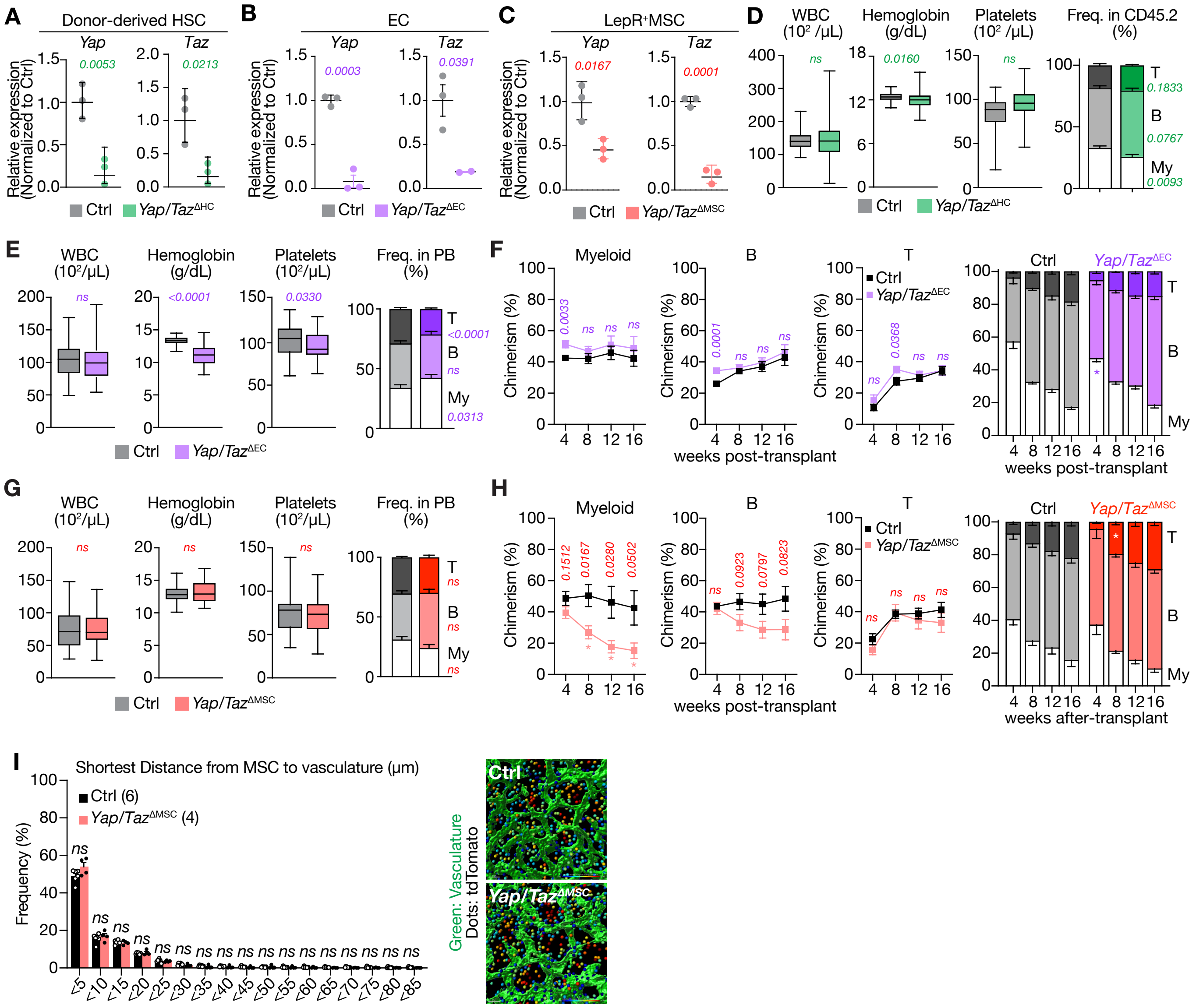

**Supplemental Figure 3 | Effects of *Yap/Taz* deletion on hematopoiesis and niche cells.** (A, B, C) mRNA expression of *Yap* and *Taz* relative to *Hprt* in BM HSCs (A), BM ECs (B), and LepR+ MSCs (C) after tamoxifen treatment in *Yap/Taz*$^{\Delta HC}$, *Yap/Taz*$^{\Delta EC}$ , and *Yap/Taz*$^{\Delta MSC}$ mice, respectively, assessed by RT-qPCR (n = 3). (D) PB cell counts and myeloid, B, and T cell frequencies in Ctrl and *Yap/Taz*$^{\Delta HC}$ mice. Ctrl (n = 33) and *Yap/Taz*$^{\Delta HC}$ (n = 33) for blood cell counts and lineage cell frequencies, respectively. (E, G) PB cell counts and frequencies of each cell lineage in *Yap/Taz*$^{\Delta EC}$ (E, Ctrl, n = 5; *Yap/Taz*$^{\Delta EC}$; n = 16) and *Yap/Taz*$^{\Delta MSC}$ (G, Ctrl, n = 22; *Yap/Taz*$^{\Delta MSC}$, n = 7) mice. (F, H) The chimerism of donor-derived PB hematopoietic cells (CD45.2/CD45.1+CD45.2) at 16 weeks post-transplantation of *Yap/Taz*$^{\Delta EC}$ (F, Ctrl, n = 18; *Yap/Taz*$^{\Delta EC}$, n = 16) and *Yap/Taz*$^{\Delta MSC}$ (H, Ctrl, n = 22; *Yap/Taz*$^{\Delta MSC}$, n = 7) BM cells along with competitor BM cells (left). The frequencies of each cell lineage in PB are shown (right). (I) The distance between Ebf3-tdTomato+ MSCs and Emcn+ sinusoidal vessels in Ctrl (n = 6) and *Yap/Taz*$^{\Delta MSC}$ BM (n = 4) (upper). Representative whole-mount BM images of Ctrl and *Yap/Taz*$^{\Delta MSC}$ femurs showing signals of Emcn (green), Ebf3-tdTomato (red), and DAPI (blue) (lower). Data are shown as mean ± SEM. P-values were calculated using two-tailed Student's *t*-test (A, B, C, D, E, F, G, H) and one-way ANOVA with Holm-Sidak post-hoc test (I). n.s., not significant.

### Supplemental Figure 4

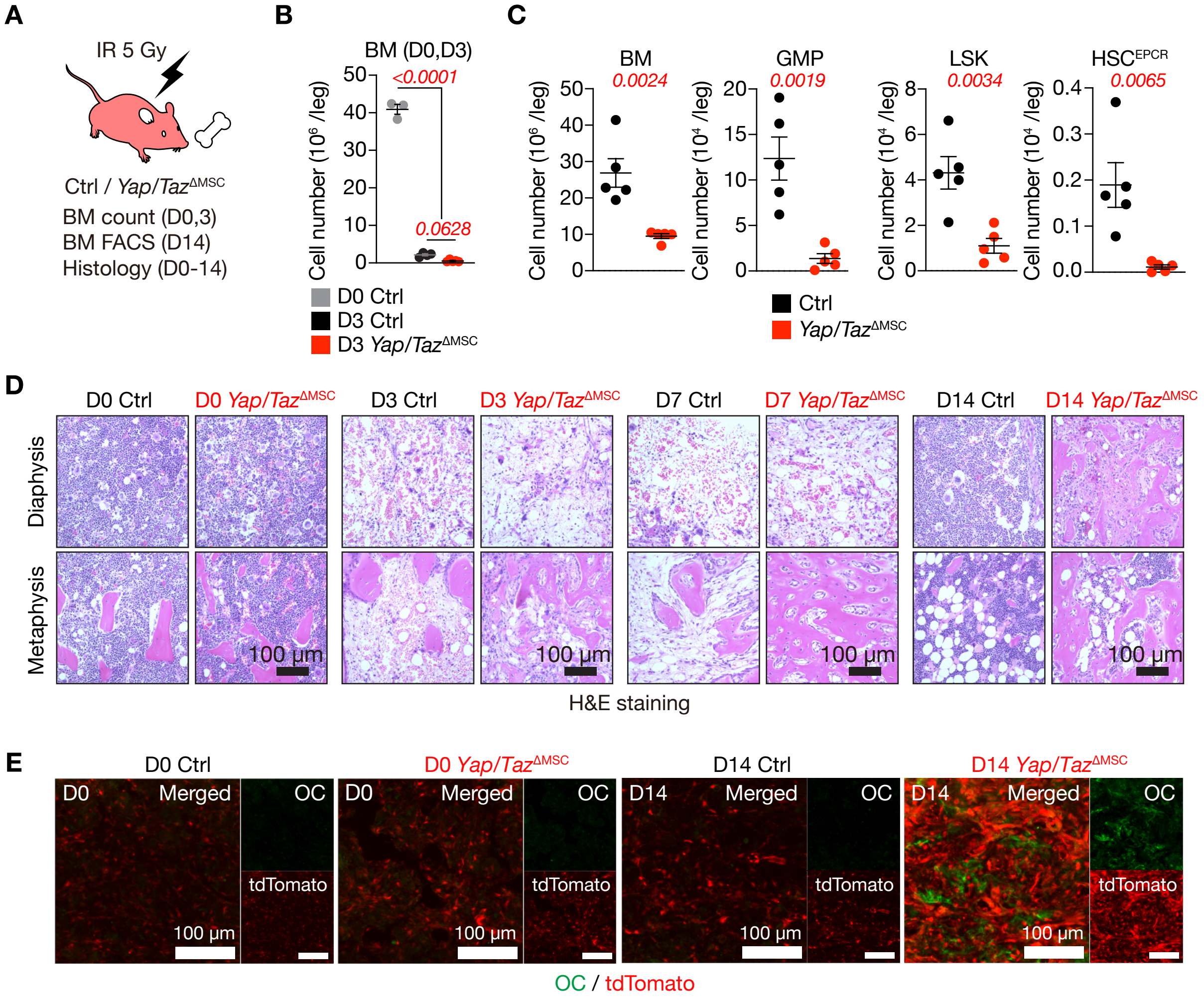


**Supplemental Figure 4 | Differentiation of *Yap/Taz*-deficient MSCs after IR.**
(A) Schematic diagram for analyzing BM in *Yap/Taz*$^{\Delta MSC}$ mice. (B) Absolute numbers of BM cells in Ctrl and *Yap/Taz*$^{\Delta MSC}$ mice before and 3 days after IR in a unilateral leg (n = 3–4). (C) Absolute cell numbers of total BM, GMP, LSK, and HSC$^{EPCR}$ populations in a unilateral leg (femur and tibia) of Ctrl and *Yap/Taz*$^{\Delta MSC}$ mice at day 14 post-IR. Data are shown as the mean ± SEM (n = 4–6). (D) H&E staining of Ctrl and *Yap/Taz*$^{\Delta MSC}$ BM sections day 0-14 after IR. Scale bars, 100 µm. (E) Representative BM images of thick femur sections from *Ebf3-tdTomato* Ctrl and *Yap/Taz*$^{\Delta MSC}$ mice after 5 Gy of IR showing Osteocalcin (OC, green) and Ebf3-tdTomato (tdTom, red). Scale bars, 100 µm. Data are shown as mean ± SEM. P-values were calculated using two-tailed Student's *t*-test (C) and one-way ANOVA with Holm-Sidak post-hoc test (B). n.s., not significant.

### Supplemental Figure 5

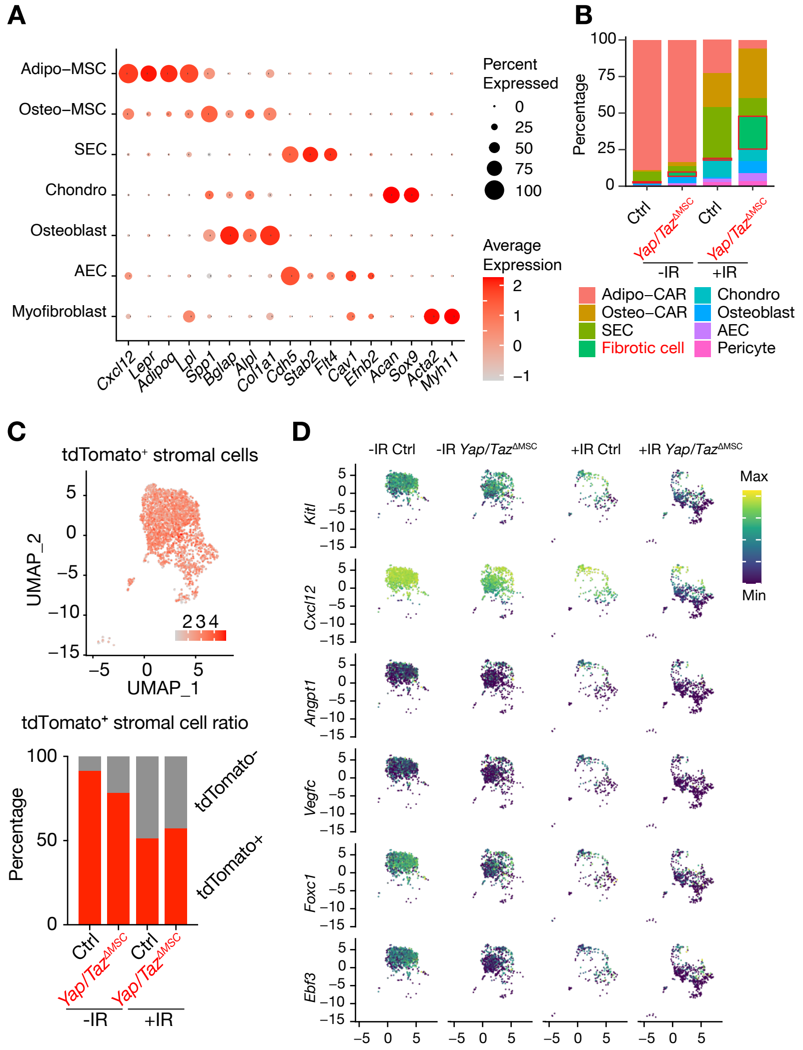


**Supplemental Figure 5 | scRNA-seq analysis of *Yap/Taz*-deficient MSCs.**
(A) Dot plots showing the expression levels of signature genes in BM ECs and stromal cells. (B) Proportional changes in ECs and stromal cells isolated from Ctrl and *Yap/Taz*$^{\Delta MSC}$ BM before and 3 days after IR. (C) UMAP illustration of *Ebf3-tdTomato*$^{+}$ Ctrl and *Yap/Taz*$^{\Delta MSC}$ stromal cells before and 3 days after IR. (upper). tdTomato+ cell proportion among stromal cells in Ctrl and *Yap/Taz*$^{\Delta MSC}$ BM before and 3 days after IR. (D) UMAP illustration showing the expression levels of key MSC genes.

### Supplemental Figure 6

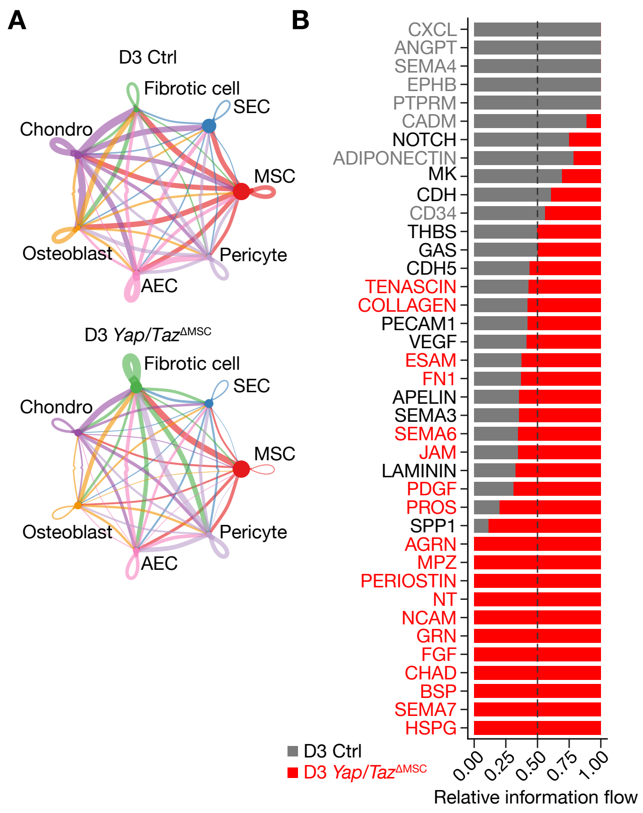


**Supplemental Figure 6 | Ligand-receptor analysis among *Yap/Taz*$^{\Delta MSC}$ BM niche cells.** (A) Circle plots showing all cell-cell interactions found in Ctrl and *Yap/Taz*$^{\Delta MSC}$ BM niche components. (B) Cell-cell communication pathways significantly enriched among niche components isolated from *Yap/Taz*$^{\Delta MSC}$ mice compared with controls. Bars represent the relative information flow for each respective signaling network, which represents relative interaction strength of the indicated ligand-receptor pairs in Ctrl (gray) and *Yap/Taz*$^{\Delta MSC}$ niche (red) cells.

### Supplemental Figure 7

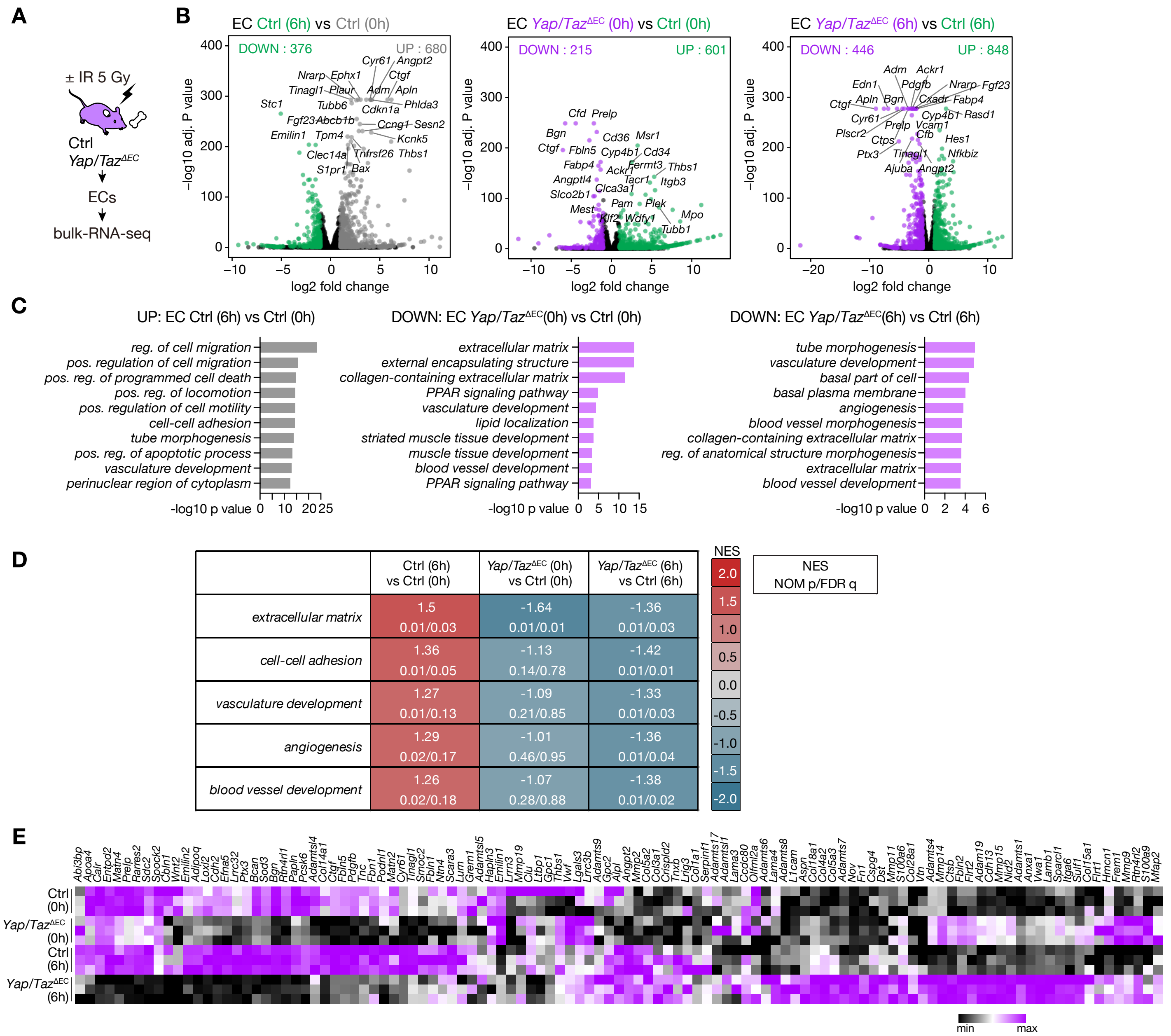


**Supplemental Figure 7 | YAP/TAZ regulate angiogenesis-related genes in sinusoidal ECs.**
(A) Schematic diagram for bulk RNA-seq in Ctrl and *Yap/Taz*$^{\Delta EC}$ ECs at steady-state or after sublethal irradiation. (B) Volcano plot showing differentially expressed genes (adjusted P < 0.05, log2 Fold Change > 2) based on bulk RNA-seq data of steady-state and sublethally irradiated Ctrl and *Yap/Taz*$^{\Delta EC}$ ECs (n = 3). Differentially expressed genes are highlighted in green, purple, and gray. (C) GO terms enriched in genes upregulated in Ctrl ECs (6 h) compared to Ctrl ECs (0 h) and downregulated in *Yap/Taz*$^{\Delta EC}$ ECs compared to Ctrl ECs (0 and 6 h). (D) Selected GSEA gene sets that were differentially enriched in each comparison group. (E) Heatmap displaying the expression levels of genes associated with angiogenesis, cell adhesion, and extracellular matrix.

### Supplemental Figure 8

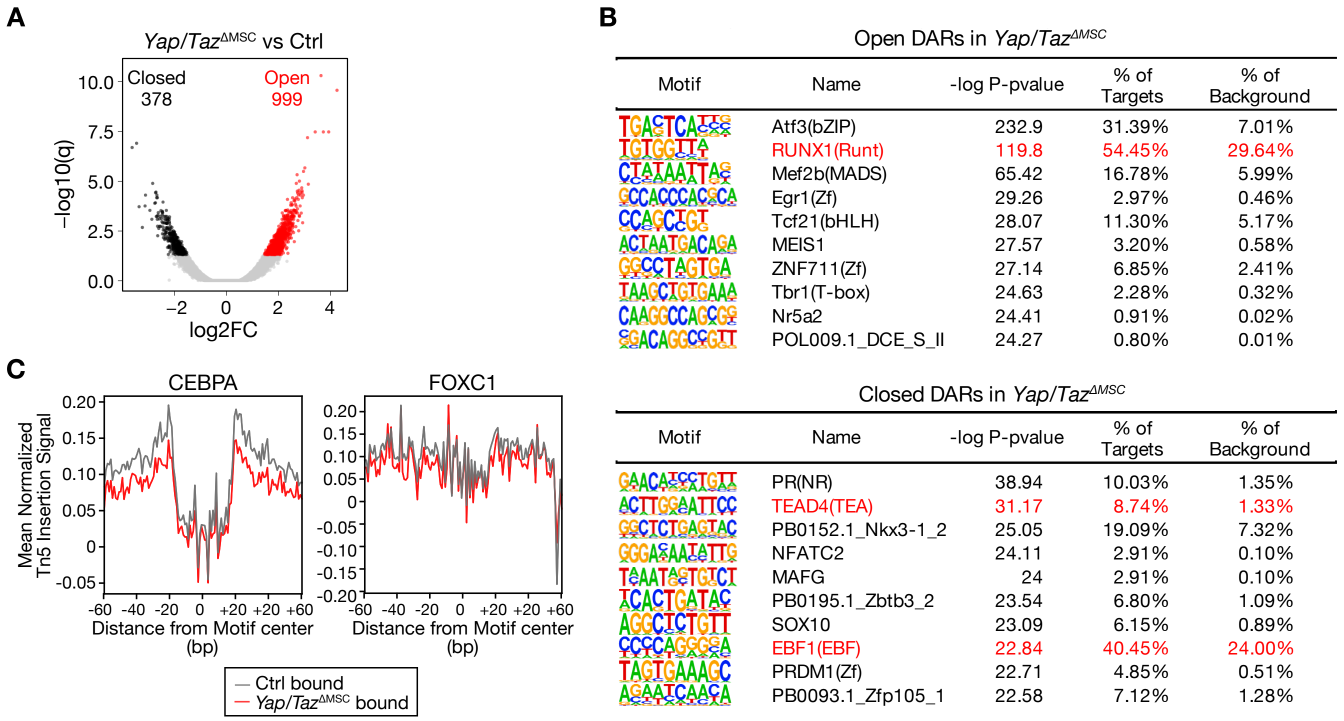


Open DARs in *Yap/Taz*$^{\Delta MSC}$

| Motif | Name | -log P-pvalue | % of Targets | % of Background |
|---|---|---|---|---|
| | Atf3(bZIP) | 232.9 | 31.39% | 7.01% |
| | RUNX1(Runt) | 119.8 | 54.45% | 29.64% |
| | Mef2b(MADS) | 65.42 | 16.78% | 5.99% |
| | Egr1(Zf) | 29.26 | 2.97% | 0.46% |
| | Tcf21(bHLH) | 28.07 | 11.30% | 5.17% |
| | MEIS1 | 27.57 | 3.20% | 0.58% |
| | ZNF711(Zf) | 27.14 | 6.85% | 2.41% |
| | Tbr1(T-box) | 24.63 | 2.28% | 0.32% |
| | Nr5a2 | 24.41 | 0.91% | 0.02% |
| | POL009.1_DCE_S_II | 24.27 | 0.80% | 0.01% |

Closed DARs in *Yap/Taz*$^{\Delta MSC}$

| Motif | Name | -log P-pvalue | % of Targets | % of Background |
|---|---|---|---|---|
| | PR(NR) | 38.94 | 10.03% | 1.35% |
| | TEAD4(TEA) | 31.17 | 8.74% | 1.33% |
| | PB0152.1_Nkx3-1_2 | 25.05 | 19.09% | 7.32% |
| | NFATC2 | 24.11 | 2.91% | 0.10% |
| | MAFG | 24 | 2.91% | 0.10% |
| | PB0195.1_Zbtb3_2 | 23.54 | 6.80% | 1.09% |
| | SOX10 | 23.09 | 6.15% | 0.89% |
| | EBF1(EBF) | 22.84 | 40.45% | 24.00% |
| | PRDM1(Zf) | 22.71 | 4.85% | 0.51% |
| | PB0093.1_Zfp105_1 | 22.58 | 7.12% | 1.28% |

**Supplemental Figure 8 | Chromatin accessibility and footprint profiles in *Yap/Taz*-deficient MSCs.**
(A) Volcano plot showing differentially accessible regions (DARs; adjusted P-value < 0.05, log2 Fold Change > 2) based on the ATAC-seq data of Ctrl and *Yap/Taz*$^{\Delta MSC}$ MSCs (n = 2). Significantly open and closed DARs are indicated in red and black, respectively. (B) Analysis of *de novo* transcription factor binding motifs in open and closed DARs in *Yap/Taz*$^{\Delta MSC}$ MSC. Motifs and log10 P-values are shown. The enrichment of each motif in DARs and background peaks in *Yap/Taz*$^{\Delta MSC}$ MSC are indicated. (C) Average normalized Tn5 insertion signal around the CEBPA and FOXC1 motifs calculated using TOBIAS.

### Supplemental Figure 9

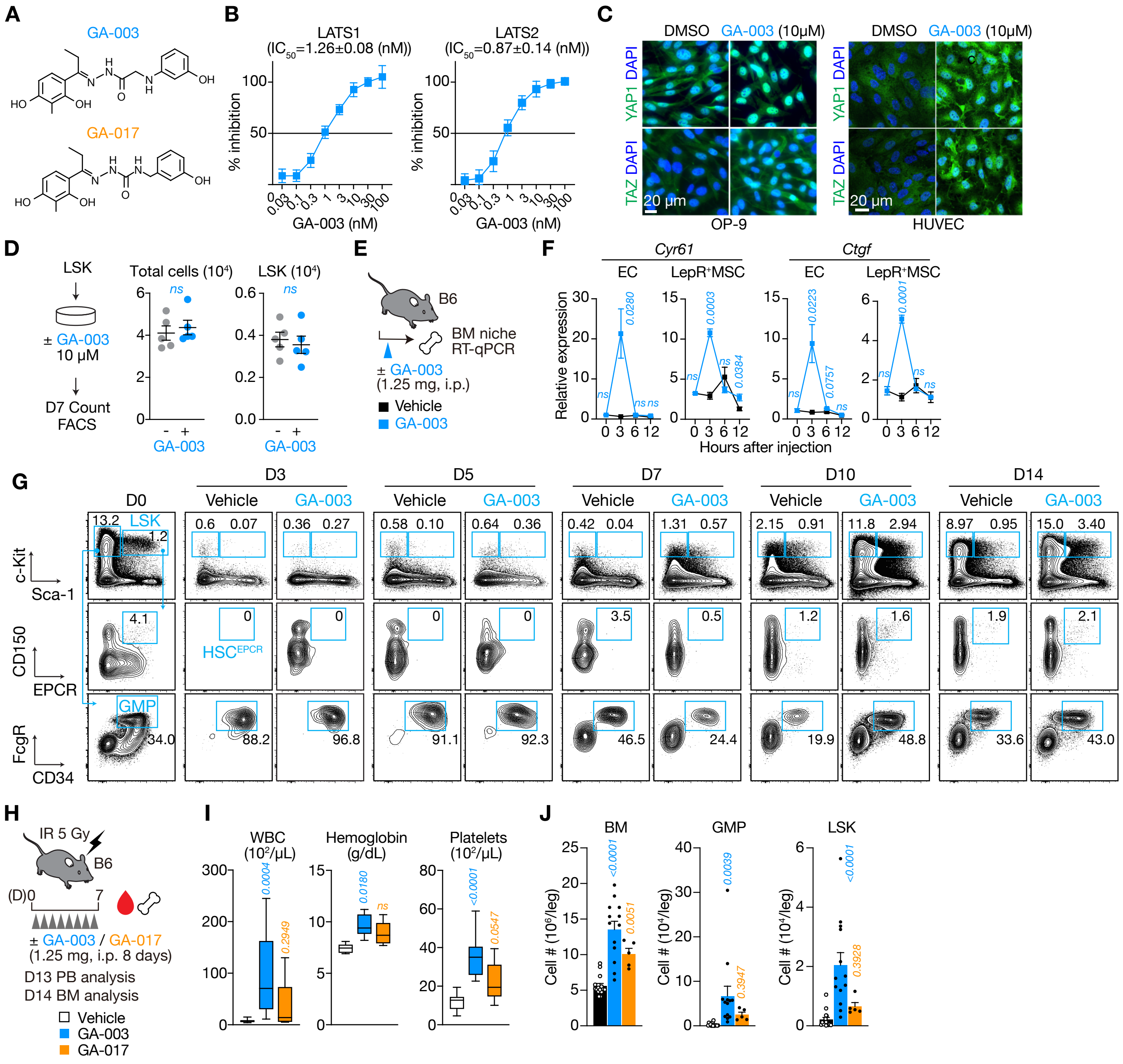

**Supplemental Figure 9 | Effects of GA-003 on hematopoietic regeneration.**
(A) Chemical structures of GA-003 (upper) and GA-017 (lower). (B) $IC_{50}$ of GA-003 against LATS1/2 kinase activity (n = 3). (C) Representative immunofluorescence images of YAP or TAZ (green) and DAPI (blue) in OP-9 cells and HUVECs 3 hours post-GA-003 treatment. (D) Growth of LSK cells in culture. The total and absolute numbers of LSK cells in culture were examined on day 7 (n = 5). (E) Experimental design to assess the efficacy of GA-003 *in vivo*. (F) mRNA expression of *Cyr61* and *Ctgf* relative to *Hprt* in BM ECs and LepR$^+$ MSCs at 0–12 hours post-GA-003 injection assessed by RT-qPCR (n = 3). (G) Representative flow cytometric profiles of BM immature cells following IR and GA-003 treatment, related to Figure 6C. (H) Experimental design of GA-003/GA-017 treatment of sublethally irradiated WT mice. (I, J) Complete blood counts (Ctrl, n = 15; GA-003, n = 13; GA-017, n = 15) on day 13 (I) and absolute cell numbers of total BM, GMP, and LSK populations (Ctrl, n = 7; GA-003, n = 10; GA-017, n = 5) (J) on day 14 post-IR. Data are shown as mean ± SEM. P-values were calculated using two-tailed Student's *t*-test (D, F) and one-way ANOVA with Holm-Sidak post-hoc test (I,J). n.s., not significant.

### Supplemental Figure 10

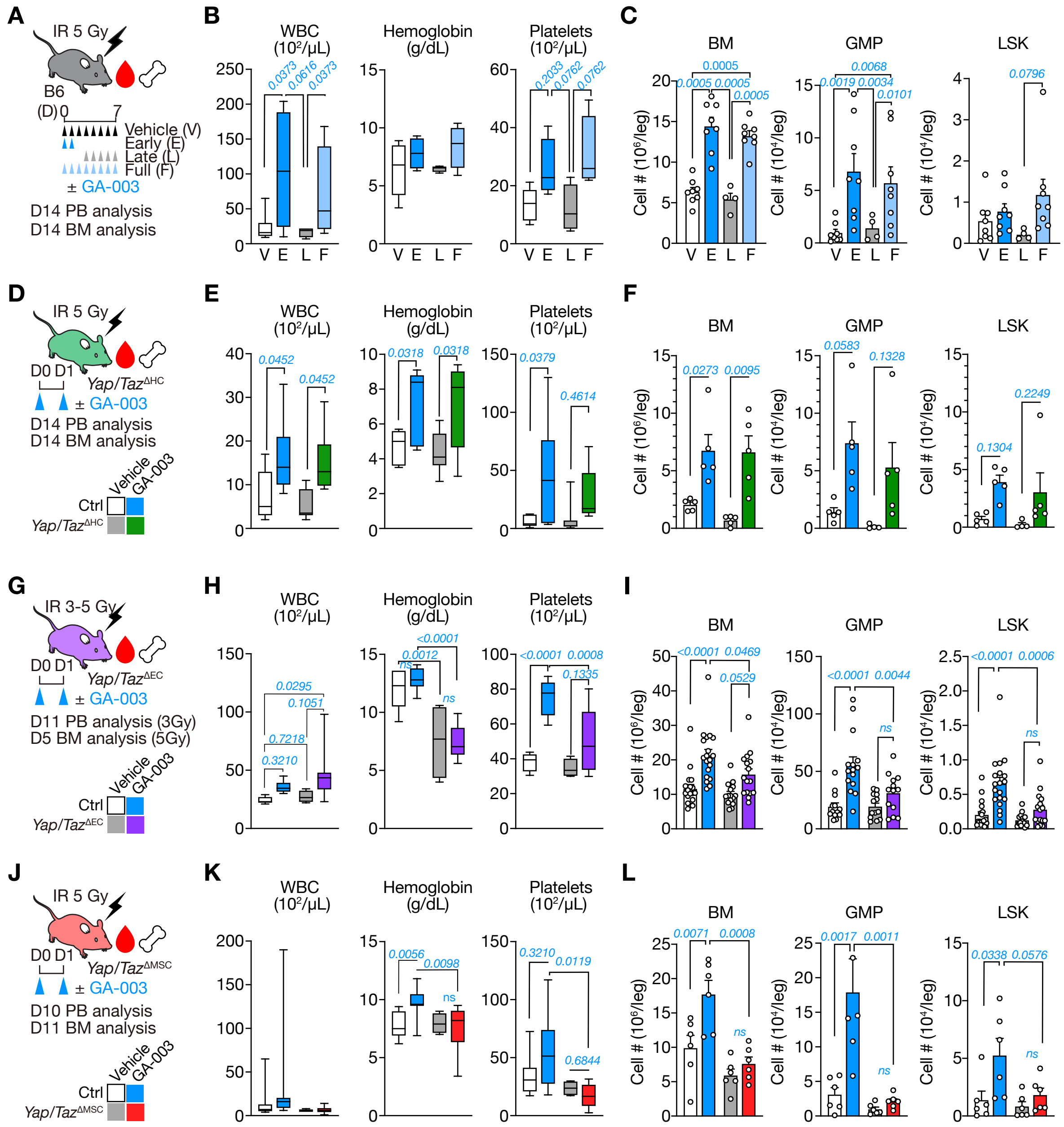


**Supplemental Figure 10 | Niche-mediated effects of GA-003 on hematopoietic regeneration.**

(A-C) Experimental design for dose de-escalation of GA-003 treatment (A). Complete blood counts (n = 4–5) (B) and absolute cell numbers of total BM, GMP and LSK populations (n = 4–8) on day 14 post-IR (C). (D-L) Experimental design of GA-003 treatment of sublethally irradiated *Yap/Taz*$^{\Delta HC}$ (D), *Yap/Taz*$^{\Delta EC}$ (G), and *Yap/Taz*$^{\Delta MSC}$ (J) mice. Complete blood counts (E, n = 6–8; H, n = 4–10; K, n = 4–10) and absolute numbers of BM, GMP, and LSK populations (F, n=4–5; I, n = 15–19; L, n = 6–7) at the indicated time points after IR. P-values were calculated using one-way ANOVA with Holm-Sidak post-hoc test. n.s., not significant.!

### Supplemental Figure 11

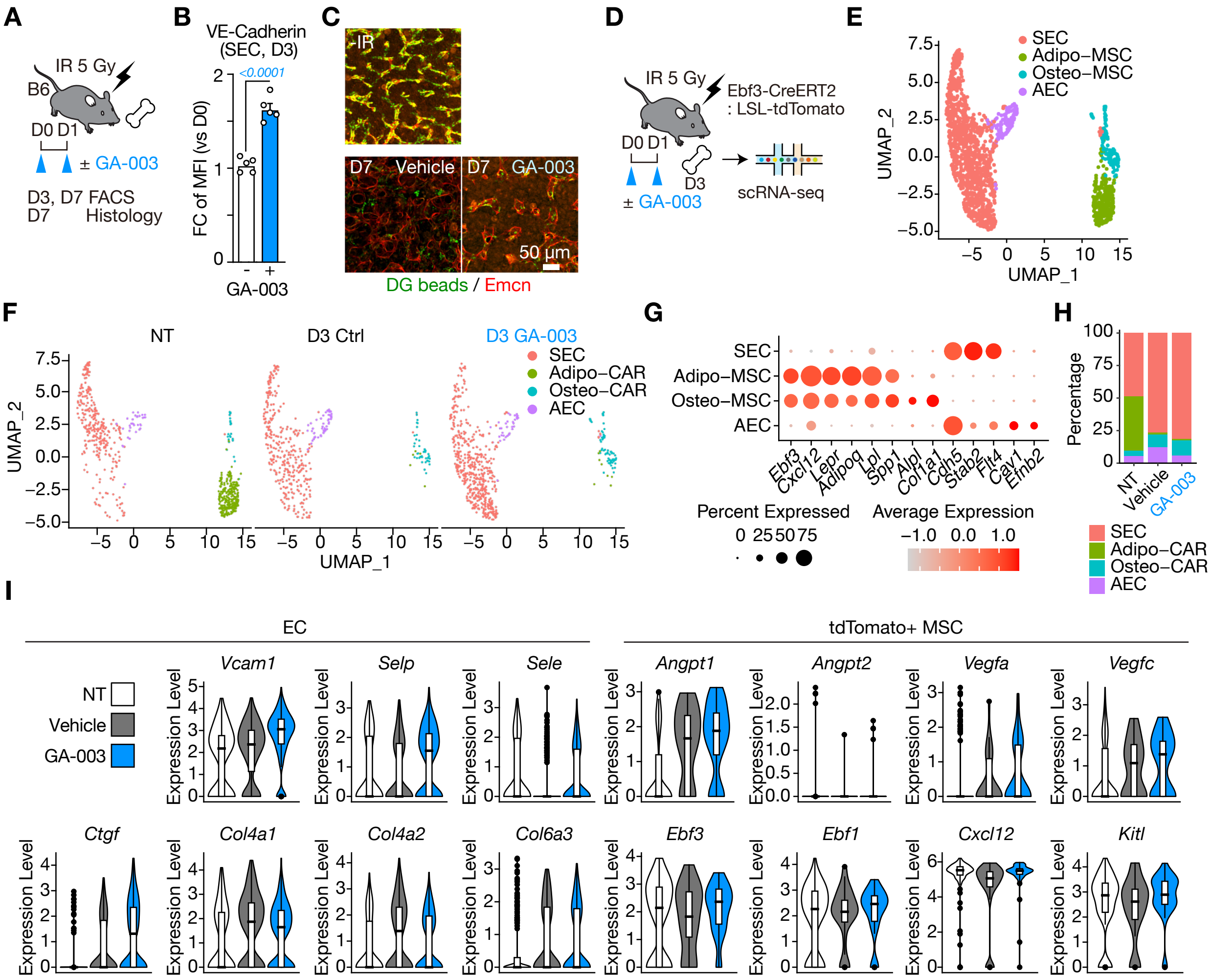


**Supplemental Figure 11 | Effects of GA-003 on BM niche components.**
(A) Experimental strategy for BM FACS and histological analysis on days 3 and 7 post-IR. (B) VE-Cadherin levels in BM ECs on day 3 (n = 5). (C) Representative immunofluorescence images of femurs from mice on day 7 post-IR. Dragon Green beads (green), and Emcn (red). (D) Experimental design for scRNA-seq analysis of BM non-hematopoietic cells isolated from *Ebf3-tdTomato* mice on day 3 post-IR and GA-003 treatment. (E-F) UMAP illustration of four cell clusters of BM niche cells based on merged scRNA-seq data of three samples (E, 1,667 cells) and UMAP plots separated into each sample (F, NT, non-treated; D3 Ctrl, irradiated and vehicle-treated; D3 GA-003, irradiated and GA-003-treated). (G) Dot plots showing the expression levels of the signature genes in BM ECs and stromal cells. (H) Proportional changes in ECs and stromal cells isolated from vehicle- or GA-003-treated *Ebf3-tdTomato* mice before and 3 days after IR. (I) Violin plot showing the expression of ECM and cell adhesion molecule genes in ECs (left) and key transcription factor and cytokine genes in *Ebf3-tdTomato*$^+$ stromal cells (right). P-values were calculated using two-tailed Student's *t*-test. n.s., not significant.
