## SupplementalText for "Niche-targeted therapy via YAP/TAZ activation enhances hematopoietic regeneration"

### Supplemental Methods

#### Mice

*Rosa26-tdTomato*[1] mice were obtained from the Jackson Laboratory. Eight-week-old NOD.Cg-Prkdc$^{scid}$Il2rg$^{tm1Wjl}$/SzJ. (NSG) mice were purchased from Central Institute for Experimental Animals (CIEA). To achieve hematopoietic-specific deletion of *Yap1/Taz* mice (*Yap/Taz*$^{\Delta HC}$), we transplanted total BM cells (5×10$^6$) of *Rosa26-CreER*$^{T2-/+}$*;Yap1*$^{fl/fl}$*;Taz*$^{fl/fl}$ mice into lethally irradiated (4.75 Gy ×2) B6-CD45.1 recipient mice (8–10-week-old). 2 months after BM transplantation, the transplanted mice were injected intraperitoneally with 100 µL of 10 mg/ml tamoxifen for five consecutive days. *Yap/Taz*$^{\Delta HC}$ mice were then analyzed 4–5 weeks after the last tamoxifen injections. To generate *Ebf3-tdTomato* (*Ebf3-CreER*$^{T2+/-}$*;LSL-tdTomato*$^{+/-}$) mice, we crossed *Ebf3-CreER*$^{T2}$ mice with *LSL-tdTomato* mice. To label *Ebf3*$^+$ MSC with tdTomato, 8–10-week-old *Ebf3-tdTomato* mice were fed with TAM food for 14 days. *LepR-Cre;Ebf1*$^{fl/+}$*;Ebf3*$^{fl/fl}$ mice were analyzed at the age of 34 weeks. Primer oligonucleotide sequences for genotyping PCR are shown in Supplemental Table 7.

#### Human cord blood cells

Human cord blood cells were provided by the Japanese Red Cross Society Kanto-Koshinetsu Cord Blood Bank. CD34$^+$ cells were obtained from cord blood mononuclear cells using Lymphoprep (STEMCELL technologies) and CD34 MicroBeads (Miltenyi Biotec, 130-046-702), then cryopreserved in CELLBANKER1 (Zenogen Pharma). For transplantation, the cells were thawed and cultured under 37℃ and 2% CO2 in StemSpan SFEM II serum-free media (STEMCELL Technologies) for 24 hours.

#### *In vivo* assays

For sublethal ionizing irradiation, mice were exposed to doses of 5.0 Gy using an X-ray irradiator (Hitachi, MSBR-1520R-4) except for the kinetic analysis of PB in *Yap/Taz*$^{\Delta EC}$ mice exposed to 3.0 Gy. For mouse cell transplantation, 8–12-week-old B6-CD45.1 recipient mice were lethally irradiated (9.5 Gy, delivered in split doses 3 hours apart) and 5×10$^6$ whole BM cells were transferred by retroorbital injection to generate *Yap/Taz*$^{\Delta HC}$ mice. For HSC transplantation, 250 donor CD45.2 HSCs (250) were transplanted with 5×10$^6$ Sca1-depleted B6-CD45.1 helper BM cells. For xenograft transplantation, 10-week-old NSG mice were exposed to a dose of 2.5 Gy and transplanted with $1.0\times10^4$ $CD34^+$ cord blood cells. For 5-FU treatment, mice were injected intraperitoneally with 150 mg/kg 5-FU (Kyowa KIRIN) dissolved in 300 μL of PBS. All recipient mice and 5-FU-treated mice were administered enrofloxacin (Elanco) in drinking water for four weeks. PB was obtained by bleeding from the tail vein and dispensed into tubes containing 0.5 μL of 0.5 M EDTA. Complete blood count (CBC) analysis was performed using a Celltac α CBC analyzer (Nihon Kohden, MEK-6558), and flow cytometry analysis was performed after RBCs were lysed in ACK lysis buffer (150 mM $NH_4Cl$, 10 mM KHCO3). BM cells were obtained by crushing leg, arm, and pelvic bones  or flushing leg bones and resuspended in 2% fetal bovine serum (FBS)/PBS (SM). After RBCs were lysed in ACK lysing buffer containing 0.02% $NaN_3$, BM cellularity was determined using a ViCELL-XR cell counter (Beckman-Coulter). The entire circulating PB cell compartment was collected by transcardiac perfusion with 20 mL of 10 mM EDTA/PBS.

**Flow cytometry**

For mouse HSC and progenitor cell sorting, BM cells were pre-enriched for c-Kit positive cells using APC-conjugated anti-c-Kit (2B8) antibody (BioLegend) and anti-APC microbeads (Miltenyi Biotec, 130-090-855). For analysis and sorting of immature hematopoietic cells, unfractionated or c-Kit-enriched BM cells were then incubated with biotinylated anti-lineage antibodies against Gr-1 (RB6-8C5), Mac1 (M1/70), Ter-119 (TER-119), CD4 (GK1.5), CD8α (53–6.7), and B220 (RA3-6B2) (BioLegend), followed by fluorophore-conjugated streptavidin and antibodies against c-Kit (2B8), Sca-1 (D7), Flk2 (A2F10), CD34 (RAM34), CD150 (TC15-12F12.2), and CD48 (HM48-1). For analysis and sorting of PB and BM mature cells, unfractionated cells were stained with antibodies against Gr-1(RB6-8C5), B220 (RA3-6B2), CD19 (6D5), Mac1 (M1/70), CD4 (GK1.5), CD8α (53–6.7), CD3e (145-2C11), Ter-119 (TER-119), CD45.1 (A20), and CD45.2 (104). For BM analysis after xenograft transplantation, cells were stained with biotinylated anti-lineage antibodies against human CD2 (RPA-2.10), CD3 (UCHT1), CD14 (61D3), CD16 (3G8), CD19 (HIB19), CD20 (2H7), CD56 (HCD56), CD66b (G10F5), and CD235a (HIR2), followed by fluorophore-conjugated streptavidin and antibodies against human CD45RA (HI100), CD34 (561), CD45 (H130), CD38 (HIT2), FLK2 (BV10A4H2), CD10 (HI10a), as well as mouse CD45 (30-F11) and Ter119 (TER-119). For analysis and sorting of BM niche cells, BM cells were flushed out from femurs and tibiae and incubated in DMEM (Sigma-Aldrich, D5796) containing 4 mg/ml collagenase type I (Worthington, LS004194) and 0.2 mg/ml DNase I (Sigma-Aldrich, D5025) 25 minutes at 37 ℃ with gentle shaking. After RBCs were removed by ACK lysis, cells were stained with antibodies recognizing LepR (BAF497), CD45 (30F11), PDGFRα (APA5), Sca-1 (D7), CD31 (390), Ter119 (TER-119), CD51 (BV13), CD105 (MJ7/18), and CD144 (TY/23). Dead cells were stained with 0.5 μg/ml propidium iodide (Sigma-Aldrich, P4170) and excluded from cell sorting and analysis. Cell sorting was performed on FACSAriaIIIu (BD Biosciences). All data were collected on FACSAriaIIIu or FACSCelesta (BD Biosciences) and analyzed with the FlowJo (BD Biosciences). Gating strategies used to identify each hematopoietic population are shown in Supplemental Figure 1.

**_In vitro_ assays**

500 mouse LSK cells were directly sorted into individual wells of 96-well round bottom plates containing 200 μL of S-Clone SF-O3 (Sanko Junyaku) supplemented with 0.2% bovine serum albumin (BSA, STEMCELL Technologies, 09300), 50 μM 2-mercaptoethanol (2-ME, Sigma-Aldrich), 100 U/ml penicillin, 100 μg/ml streptomycin, 292 μg/ml L-Glutamine (1× PSG, Gibco, 10378016), 50 ng/ml mouse SCF (BioLegend, 579706) and 50 ng/ml human TPO (BioLegend, 763706). Cells were cultured for 7 days at 37℃ in a 5% CO2 incubator and counted using a Vi-CELL-XR.

**CFU-F assay**

BM cells flushed out from the femur and tibia were digested with 4 mg/ml collagenase type I as described above. After RBCs were removed by ACK lysis, one-tenth of the cells were seeded into individual wells of a 6-well plate according to MesenCult expansion kit protocol (STEMCELL Technologies, 05513). Cells were cultured at 37℃ under hypoxic conditions (5% $O_2$ and 5% $CO_2$) for 7 days and stained with May-Grunwald (Sigma-Aldrich) and Giemsa (Sigma-Aldrich) solutions. Colony-forming units and cells per colony were counted visually under microscope.

**Immunostaining of OP-9 cells and HUVECs**

OP-9 cells were maintained in α-MEM supplemented with 2-ME, 1× PSG, and 20% FBS. HUVECs were maintained in Endothelial Cell Growth Medium (Promo Cell, C-22010) supplemented with ×42 supplement mix (Promo Cell, C-22010) and 1× PSG. For immunostaining, $1\times10^5$ OP-9 cells or HUVECs were seeded into individual wells of 8-well chamber glass slides (SPL, SPL-30108), precoated with gelatin (Sigma-Aldrich, G1890), and cultured for 96 hours until confluent at 37 °C in a 5 % $CO_2$ incubator. Cells were fixed in 4% PFA/PBS (Thermo Fisher, 28906), permeabilized in 0.1% Triton X-100/PBS (Sigma-Aldrich, T9284), blocked in 3% BSA/PBS (Sigma-Aldrich, A9647), and stained with rabbit anti-YAP (Cell Signaling, 14074S, 1:400) and rabbit anti-TAZ (Cell Signaling, 83669S, 1:400) antibodies overnight at 4℃. Following three PBS washes, cells were stained with fluorophore-conjugated donkey anti-rabbit IgG (Biotium, 20015, 1:1000) antibody for 2 hours at room temperature (RT). Cells were then counterstained in 1 μg/mL DAPI/PBS for 5 min at RT and mounted with ProLong Glass Antifade Mountant (Thermo Scientific, P36982). Images were captured on a Dragonfly confocal microscope (Andor) with a 40× objective or a BZ-9000 fluorescence microscope (KEYENCE) with a 40× objective. Images were processed with Imaris (Bitplane) and Fiji.

**BM histological staining**

Upon isolation, mouse femurs were fixed in 10% formalin neutral buffer solution (Wako, 062-01661) at 4°C for 36 hours with gentle agitation, decalcified in 0.5 M EDTA (pH 7.5) (Dojindo, 345-01865) for 7 days, and embedded in paraffin. For H&E staining, the BM section was stained in hematoxylin (Muto pure chemicals, 3000-2) and eosin (Muto pure chemicals, 3204-2) solutions and mounted with malinol (Muto pure chemicals, 2040-1). For reticulin silver staining, the BM section was stained in ammoniacal silver solution (Muto pure Chemicals, 40411) according to the manufacturer's instructions. The stained BM was imaged on a BX53 microscope (Olympus) with a 2× (EVIDENT, PlanApo N 2x/0.08NA) or 40× (EVIDENT, MPlanFL N 40×/0.75NA) objective.

**BM immunostaining**

For *in vivo* staining of BM vessels, mice were injected retro-orbitally with a cocktail of CD144-Alexa Fluor 647 (BioLegend, 138006) and CD31-Alexa Fluor 647 (BioLegend, 102416) antibodies 10 minutes before euthanasia. For vascular permeability analysis, mice were injected retro-orbitally with 2.5 μl/g body weight of Dragon Green beads (DGB; Bangs Laboratories, FSDG001) 10 min before euthanasia. Upon isolation, femurs were fixed in ice-cold 2% PFA/PBS for 16 hours with gentle agitation, decalcified in 0.5 M EDTA (pH 7.5) for 7 days, sequentially immersed overnight in 15% and 30% sucrose (Sigma-Aldrich, 28-0010-5), embedded in OCT (Sakura, 4583), and snap frozen. For staining of BM sections, frozen femurs were cryosectioned at a thickness of either 10 μm using Kawamoto's film method[2] or 80 μm using a cryostat (Cryostar NX70, Epredia). Sections were transferred onto glass slides, blocked for 1 hour at RT in TBS containing 10% normal donkey serum (Jackson Immuno Research, 017-000-121), with 0.05% Tween-20 (BIO-RAD, 170-6531) or 0.3% Triton X-100 (Sigma-Aldrich, X100) added for intracellular staining, and stained with rat anti-endomucin (Santa Cruz, sc-65495, 1:400), rat anti-osteocalcin (Takara, M188, 1:200), rabbit anti-YAP (Cell Signaling, 14074S, 1:400), and rabbit anti-TAZ (Cell Signaling, 83669S, 1:400) antibodies overnight at 4℃. Sections were washed in TBS three times and stained with fluorophore-conjugated donkey anti-rat IgG (Thermo Scientific, SA5-10029, 1 : 400) and donkey anti-rabbit IgG (Biotium, 20015, 1 : 400) antibodies for 3 hours at RT. Finally, sections were counterstained in 1 μg/ml DAPI/TBS for 10 minutes at RT and mounted with ProLong Glass Antifade Mountant (Thermo Scientific, P36980). For whole-mount BM staining, femurs were fixed and embedded as described above, trimmed by a cryostat until the bone cavity was exposed, and washed in TBS three times for 5 minutes at RT to remove OCT. Femurs were then blocked for 2 hours at RT in TBS containing 10% normal donkey serum, 0.05% Tween-20, and 20% DMSO (WAKO, 043-07216) and stained with rat anti-endomucin (Santa Cruz, sc-65495) antibody over two nights at 4℃. Femurs were carefully washed in TBS three times, stained with donkey anti-rat IgG (Thermo Scientific, SA5-10029, 1:200) antibody overnight at 4℃, and counterstained in 1 μg/ml DAPI/TBS for 30 minutes at RT. To enhance tissue transparency, femurs were cleared in 2,2-Thiodiethanol (TDE) (Sigma-Aldrich, 166782) as described previously[3]. All images were captured on a Dragonfly spinning disk confocal microscope (Andor) with a 10× (Nikon Plan Apo 10×/0.45 DIC N1 Lambda) or 20× (Nikon Plan Apo 20×/0.75 DIC M/N2) objective and processed using Imaris10 (Bitplane) or Fiji. Vascular area was calculated using Fiji's "analyze particles" function after the endomucin signal was binarized with Otsu's algorithm and the pixels within the vessels were filled in. For whole-mount BM staining, maximum intensity projections of 100 μm z-stack images are displayed unless otherwise noted.

**Chemical synthesis and *in vivo* and *in vitro* treatment of GA-003**

GA-003 [(E)-N'-(1-(2,4-dihydroxy-3-methylphenyl)propylidene)-2-((3-

hydroxyphenyl)amino)acetohydrazide] was synthesized using standard synthetic procedures as reported previously[4]. Briefly, 2-((3-hydroxyphenyl)amino)acetohydrazide (130 mg, 0.72 mmol) and 1-(2,4-dihydroxy-3-methylphenyl)propan-1-one (100 mg, 0.55 mmol) were dissolved in 1.1 mL of DMSO, and the mixture was stirred at 100°C for 21 hr. The solution was allowed to cool, and the reaction solution was directly purified by moderate-pressure silica gel column chromatography (silica gel 10 g, ethyl acetate/methylene chloride = 5/95–50/50). The obtained solid was washed with water and isopropyl ether and dried under reduced pressure to give GA-003 (95.9 mg, 0.279 mmol, yield 51%) as a white solid. For *in vivo* treatment, GA-003 powder was ground with a mortar and pestle to make the particles homogeneous and then suspended in 0.5w/v% methylcellulose (WAKO, 133-17815) to make 6.25 mg/mL GA-003 solution. 200 µL of GA-003 solution was injected into a mouse every 24 hours. For treatment of irradiated mice, GA-003 was first administered 30 minutes before exposure to IR. For *in vitro* treatment, GA-003 powder was dissolved in DMSO (WAKO, 031-24051) and then mixed with the medium to a final concentration of 0.1% DMSO.

**Kinase assay**

Kinase assay was performed using a Fluorospark® Kinase/ADP Multi-Assay kit (WAKO, 291-77401) at a final volume of 5 µL/well as described previously[4].

**Quantitative RT-PCR**

Total RNA was extracted using RNeasy Micro Plus Kit (QIAGEN, 74134) and transcribed using SuperScript IV First-Strand Synthesis System (Invitrogen, 18091050) and oligo-dT primers. Real-time quantitative PCR was performed on a StepOnePlus Real-Time PCR System (Life Technologies) using TB Green Premix Ex Taq II (Takara Bio, RR820). All data are presented as relative expression levels normalized to *Hprt* expression. The primer sequences used are shown in Supplementary Table 8.

**Bulk RNA sequencing**

Total RNA was extracted from 10,000 cells using RNeasy Plus Micro Kit, and cDNA was synthesized using SMART-Seq v4 Ultra Low Input RNA Kit for Sequencing (Takara, 634888) according to the manufacturer's instructions. The double-stranded cDNA was fragmented using S220 Focused-ultrasonicator (Covaris), then cDNA libraries were generated using NEBNext Ultra DNA Library Prep Kit (New England BioLabs, E7370L) according to the manufacturer's instructions. Sequencing was performed using NextSeq 500 (Illumina) with a single-read sequencing length of 70 bp. Sequencing quality control was performed using FastQC v0.12.1 (Babraham Bioinformatics). TopHat 2.0.13 (with default parameters) was used to map the reads to the reference genome (UCSC/mm10) with annotation data from iGenomes (Illumina), and transcript per million (TPM) for each gene was quantified with Cuffdiff (Cufflinks 2.2.1; with default parameters)[5]. Differentially expressed genes were identified using adjusted q-value < 0.05 and log2 fold change < –1. Gene ontology (GO) pathway analysis was performed using g:Profiler[6]. Gene set enrichment analyses (GSEA) were performed using C3 and C5 in MSigDB[7]. Differential gene expression was determined using DEseq2 1.42.1[8]. Heatmaps showing Z-score of TPM were generated with hierarchical clustering of samples with Morpheus (Broad Institute).

**Single-cell RNA sequencing**

Non-hematopoietic cells (CD45$^{-}$ Ter119$^{-}$) were collected from collagenase I-treated BM as described above. mRNA from each sample was isolated, and libraries were prepared using the Chromium Next GEM Single-Cell 3′ Reagent Kits v3.1 (10X Genomics, PN-1000128). Sequencing was performed using NovaSeq 6000 (Illumina) with a pair-read sequencing length of 90 bp. Raw data files (base call files) were demultiplexed into fastq files using Cell Ranger with the mkfastq command. The cellranger count command was used for feature and barcode counting, utilizing the "refdata-gex-mm10-2020-A" reference. We recovered 4,902 cells from control (IR–), 5,489 cells from *Yap/Taz*$^{\Delta MSC}$ mice (IR–), 2,846 cells from control (IR+), 2,452 cells from *Yap/Taz*$^{\Delta MSC}$ (IR+), 2,906 cells from non-treated mice, 997 cells from vehicle-treated mice, and 1,258 cells from GA-003-treated mice after filtering. Subsequent analyses were conducted using Seurat 5.1.0. Quality filtering for each feature and cell was performed based on the following criteria: minimum cells = 3, minimum features = 200, nFeature_RNA > 200, nFeature_RNA < 2500, and percent mitochondrial genes < 5%. Feature counts were log-normalized using the NormalizeData function. Two thousand highly variable features were selected for principal component analysis (PCA). PCs 1–10 were utilized for UMAP and graph-based clustering with the functions FindNeighbors (object, reduction = "pca", dims = 1:20) and FindClusters (object, resolution = 0.5). Cell-cell communication analysis was conducted using CellChat 1.6.1. Default parameter settings were used throughout, and a truncated mean approach with a minimum of ten cells was applied for signaling inference.

**ATAC sequencing**

CD45$^{-}$ Ter119$^{-}$ CD31$^{-}$ Ebf3-tdTomato$^{+}$ cells (1.0–2.0×$10^4$ cells) were lysed in cold lysis buffer (10 mM Tris-HCl pH 7.4, 10 mM NaCl, 3 mM $MgCl_2$, 0.1% IGEPAL CA-630) on ice for 10 min. After centrifugation, nuclei pellets were resuspended in 50 μL of transposase reaction mix containing 25 μL Tagment DNA buffer (Illumina), 2.5 μL Tagment DNA enzyme (Illumina), and 22.5 μL water, incubated at 37°C for 35 minutes, and purified with a MinElute PCR Purification Kit (QIAGEN). After the optimization of PCR cycle number using SYBER Green I Nucleic Acid gel Stain (Takara Bio), transposed fragments were amplified (17 cycles) using NEBNext High Fidelity 2×PCR Master mix and index primers. Amplified libraries were purified with a MinElute PCR Purification Kit (QIAGEN), sized selected with SPRIselect (BECKMAN COULTER) (Ratios (Left-Right): 0.5-1.3), and sequencing with NovaSeq 6000 (Illumina) with a pair-read sequencing length of 60 bp. Bowtie2 2.2.6 (with default parameters) was used to map reads to the reference genome (UCSC/mm10) with annotation data from iGenomes (Illumina). Reads mapped to mitochondria were removed. To ensure even processing, reads were randomly downsampled from each sample to adjust to the smallest read number of samples. MACS 2.1.1 (with default parameters) was used to call peaks in downsampled reads. The catalogue of all peaks called in any samples was produced by merging all called peaks that overlapped by at least one base pair using bedtools. As a result, a total of 331253 merged peaks were detected and used as a map file for downstream processing. The MACS bdgcmp function was used to compute the fold enrichment over the background for all populations, and the bedtools map function was used to count fragments in the catalogue in each population. Fragment counts at each site in the catalogue were quantile normalized between samples using the PreprocessCore package in R 3.3.2. We used the HOMER package with command annotatePeaks.pl using default parameters to annotate regions with promoter and distal labels and the nearest gene, and with command findMotifsGenome.pl using default parameters to identify enriched motifs, and the catalogue of all called peaks as a background. Footprint analysis was conducted using TOBIAS (v0.14.0) with default parameters. For global footprinting, all sites classified as “bound” for EBF1, EBF3, FOXC1, CEBPA, TEAD1–4 were used as input, as defined by the default settings of the BINDetect function.

**ELISA**
For collecting BM fluids, the four long bones (two femurs and two tibiae) of each mouse were flushed out with the same 200 μL of PBS and spun at 500 × g for 5 minutes to remove BM cells. Supernatants were further clarified by spinning down at 25,000 rpm for 10 minutes, and samples were subsequently stored at –80°C until use. For cytokine measurement, 100 μL of 4×-diluted samples were analyzed with ELISA kits for ANGPT1 (LS Bio, LS-F2956-1), VEGF-C (CUSABIO, CSB-E07361M), CXCL12 (R&D systems, DY460), and SCF (R&D systems, DY455) according to the manufacturer’s protocols. Optical density was determined using a CLARIOstar Plus plate reader (BMG LABTECH, 430-502-14TA).

**Quantification and statistical analysis**
Statistical tests were performed using Prism 10 (GraphPad). Statistical significance was assessed by two-tailed Student’s *t*-test and one-way ANOVA with Holm-Sidak post-hoc test. Data are shown as mean ± standard error (SEM). ns: not significant.

**Supplemental Figure Tables**

Supplemental Table 1. DEGs in bulk RNA-seq analysis, related to Figure 1. (Log2FC ≤ -1, p_val_adj < 0.05)

Supplemental Table 2. DEGs in bulk RNA-seq analysis, related to Supplemental Figure 7. (Log2FC ≤ -1, p_val_adj < 0.05)

Supplemental Table 3. Gene enrichment analysis of DEGs in ECs

Supplemental Table 4. DARs in tdTomato-MSCs: *Yap/Taz*$^{\Delta MSC}$ vs Ctrl

Supplemental Table 5. HOMER motif analysis in open and closed DARs

Supplemental Table 6. Footprint analysis and TFBS data in biding regions of EBF1, EBF3, FOXC1, CEBPA, TEAD1, TEAD2, TEAD3, and TEAD4

Supplemental Table 7. List of oligonucleotides for genomic PCR

Supplemental Table 8. List of oligonucleotides for qPCR
